# Pericyte ablation causes hypoactivity and reactive gliosis in adult mice

**DOI:** 10.1101/2023.10.26.564269

**Authors:** Jake M. Cashion, Lachlan S. Brown, Gary P. Morris, Alastair J. Fortune, Jo-Maree Courtney, Carlie L. Cullen, Kaylene M. Young, Brad A. Sutherland

**Affiliations:** Tasmanian School of Medicine, College of Health and Medicine, University of Tasmania, Hobart, Tasmania, Australia; Menzies Institute for Medical Research, University of Tasmania, Hobart, Tasmania, Australia, 7000

**Keywords:** pericyte, ablation, fatigue, gliosis, blood-brain barrier

## Abstract

Pericytes are contractile cells that enwrap capillaries allowing them to control blood flow, maintain the blood-brain barrier and regulate immune cell trafficking in the CNS. Pericytes are lost or become dysfunctional in neurodegenerative diseases such as Alzheimer’s disease, stroke, and multiple sclerosis, but their role in health and disease is poorly understood. Our aim was to evaluate blood-brain barrier integrity and glial reactivity, and to assess behavioural phenotypes that emerge following pericyte ablation in adult mice. The delivery of tamoxifen to *PDGFRβ-CreER^T2^ :: Rosa26-DTA* transgenic mice produced a dose-dependent ablation of pericytes. A single low dose of tamoxifen ablated approximately half of all brain pericytes, and two consecutive daily high doses ablated more than 80% of brain pericytes. To determine whether pericyte ablation could induce a behavioural phenotype, we assessed patterns of voluntary movement, as well as balance and coordination using the open field and beam walk tasks. Mice with ∼50% pericyte loss travelled half the distance and spent half as much time moving in the open field as control mice. Mice with more than 80% pericyte ablation also slipped more frequently in the beam walk task than control mice. In brain cryosections from pericyte-ablated mice, blood vessel structure was unchanged, but lumen area was increased. Pericyte-ablated mice also experienced blood-brain barrier leakage, hypoxia and increased microgliosis and astrogliosis compared to control mice. Our results highlight the importance of pericytes for brain health, as pericyte loss can directly drive brain injury and behavioural alterations in mice.

## Introduction

The neurovascular unit is a multicellular structure that efficiently distributes cerebral blood flow through the brain’s capillary bed in response to neuronal demand (Cashion et al., 2023). Pericytes are a key cellular component of the neurovascular unit. Pericytes are contractile cells that regulate cerebral blood flow (Hall et al., 2014, Gonzales et al., 2020, Attwell et al., 2016), maintain the blood-brain barrier (BBB) (Armulik et al., 2010, Daneman et al., 2010), and contribute to neuroinflammatory processes by secreting and responding to inflammatory molecules, presenting antigen and phagocytosis (Torok et al., 2021, Rustenhoven et al., 2017). Pericyte dysfunction is a feature of many neurodegenerative diseases including stroke, Alzheimer’s disease, and multiple sclerosis, but it remains poorly understood (Brown et al., 2019, Cashion et al., 2023).

Cell-specific *in vivo* ablation models are powerful tools for investigating cell function and several inducible pericyte ablation models were recently used to evaluate the effect of pericyte loss in the brain (Nikolakopoulou et al., 2019, Kisler et al., 2020, Choe et al., 2022), retina (Eilken et al., 2017, Park et al., 2017), ear (Zhang et al., 2021) and heart (Cornuault et al., 2023). Within the CNS, pericyte ablation: reduced cerebral blood flow; led to neuron loss and diminished long-term memory recall performance (Nikolakopoulou et al., 2019); caused neurovascular uncoupling (Kisler et al., 2020); decreased the extent of the endothelial glycocalyx, which led to increased leukocyte capillary stalling (Choe et al., 2022); increased BBB leakage (Nikolakopoulou et al., 2019) and impaired retinal endothelial cell sprouting (Eilken et al., 2017, Park et al., 2017). However, the impact of pericyte ablation on mouse behaviour, particularly motor function, and the influence of pericyte loss on brain glia, particularly microglia, astrocytes, and oligodendrocytes, remains largely uncharacterised.

Here, we show that we can specifically ablate brain pericytes using a *PDGFRβ-CreER^T2^:: Rosa26-DTA* mouse line. Behavioural experiments show that pericyte ablation rapidly leads to reduced levels of activity and impaired motor coordination in young adult mice, without altering muscle strength. Pericyte ablation did not alter vessel structure but increased vascular expression of intracellular adhesion molecule 1 (ICAM-1) and impeded microvascular function, evidenced by increased vessel lumen area, BBB leakage and a larger number of hypoxic neurons. Pericyte ablation also triggered widespread microgliosis and astrogliosis. We conclude that pericyte ablation can directly trigger pathological changes in the brain including injury to blood vessels and the BBB, gliosis, neuroinflammation and hypoxia. The physiological changes that pericyte loss induced in young adult mice may directly contribute to the concomitant hypoactivity they experienced, which closely resembled fatigue, as well as the impaired motor coordination, and highlight the capacity for pericyte dysfunction to directly drive elements of pathophysiology experienced by people with neurological disease.

## Methods

### Transgenic mice

All animal procedures were approved by the Animal Ethics Committee, University of Tasmania (A0018606) and conformed with the Australian National Health and Medical Research Council (NHMRC) Code of Practice for the Care and Use of Animals for Scientific Purposes, 2013 (eighth edition). All results are reported in accordance with the ARRIVE guidelines (Percie Du Sert et al., 2020). Transgenic mice used for this study include *PDGFRβ-CreER^T2^* (The Jackson Laboratory, cat. #029684), *Rosa26-YFP* (The Jackson Laboratory, cat. #006148) and *Rosa26-DTA* (The Jackson Laboratory, cat. #009669) and *NG2DsRed* (The Jackson Laboratory, cat. #008241) strains. All mice were maintained on a C57BL/6J background and adult (10–50-week-old) male and female mice were used for experiments. Mice were group housed in Optimice® cages on a 12h light/dark cycle (lights on 7 A.M. to 7 P.M.) with *ad libitum* access to standard chow (Barastoc) and water.

### Tamoxifen administration and health evaluation

Tamoxifen (Tx, Sigma-Aldrich, cat. #T5648-5G) was reconstituted at 40mg/ml in corn oil (Sigma-Aldrich, cat. #C8267-500ML) and sonicated for ≥1 h until dissolved. Mice received up to 4 consecutive daily doses of Tx (100-300mg/kg as specified) by oral gavage. From day 1 of tamoxifen administration, mice were weighed and monitored daily (health score in Supplementary Table 1).

### Behavioural analyses

Prior to commencement of behavioural assessments, mice were handled for 3-5 min per day for 3 consecutive days, to reduce the stress caused by handling on the day of the study.

#### Open field

The open field task was used to evaluate locomotion (Foster et al., 2021, Tatem et al., 2014). Mice were acclimatized to the sound-proof assessment room under sodium lighting for 30 min before being placed in the open field square plexiglass arena (30cm^2^, with 20cm high walls) with low lighting (100 lux). Individual mice were habituated to the arena for 10 min on the day prior to testing. On the day of testing, individual mice were placed in the arena and allowed to freely explore for 10 min before being returned to their home cage. Mice were video recorded and their movement tracked with EthoVision XT (Noldus, v11.5, activity tracked using v17.0) with the following parameters extracted: distance travelled [as per Ferreira et al. (2021)]; time spent moving; time spent stationary; maximum acceleration during movement and activity state (low, moderate, high). Temporal changes in pixel contrast were thresholded and used as a proxy for activity. The open field arena was thoroughly cleaned with F10 SC veterinary disinfectant between mice.

#### Grip strength

A Grip Strength Meter (Melquest, cat. #GPM-100) was used to estimate muscular strength. Each mouse was weighed, positioned ready to grasp the grid mounted on the force gauge, and the equipment reset to 0 gf after stabilization. Each mouse was allowed to grasp the grid with all four paws, pulled back over the grid, and the maximum tension recorded. A test was repeated if all four paws failed to successfully grasp the grid or the mouse bit or knocked the grid. Five successful trials were recorded per mouse. We calculated force (gf) relative to body weight (g) for each mouse by dividing the mean of five successful tests by body weight.

#### Beam walk

Mouse motor coordination was assessed using data obtained from the beam walk task, conducted in a sound-proof room with fluorescent lighting as previously described (Cullen et al., 2021). Mice were acclimatized to the assessment room for 30 min prior to testing. Individual mice were placed at the end of a 10mm diameter, 1000mm long piece of wooden dowel (beam), at the opposite end to their home cage bedding. Mice were trained to cross the beam by gently supporting their tail to direct initial crossings. In subsequent test sessions, mice were recorded with an action camera (Kaiser Bass x90) as they crossed the beam. Total steps and time taken to cross the beam, as well as the number of foot slips i.e. instances when the hind paw fell below the lower limit of the beam, were manually quantified by a researcher blind to genotype and mouse identification number.

### Detection of hypoxia

Mice received 60mg/kg pimonidazole (Hypoxyprobe-1, cat. # HP2-100Kit) in 0.9% saline i.p. and were perfusion fixed 90 min later (see below). Pimonidazole was detected in brain sections by immunohistochemistry as per the manufacturer’s instructions.

### Tissue acquisition and processing

Adult male and female mice received 150mg/kg sodium pentobarbitone (Virbac, cat. #1P4291-1) i.p. prior to being transcardially perfused at 10 ml/min with PBS for 1 min, which was exchanged for 4% (w/v) paraformaldehyde (PFA, pH 7.4) in PBS for 1 min. For vessel gel casting, mice were then perfused with prewarmed (37°C) FITC-albumin gel [2.5% w/v gelatin (Sigma-Aldrich, cat. #G1890-1KG), 0.05% w/v FITC-albumin (Sigma-Aldrich, cat. #A9771-1G) in PBS] for 1 min and placed on ice for 90 min to allow the gelatin to set. Whole brains and spinal cords were removed, postfixed in 4% PFA for 1.5h at room temperature (∼21°C), and cryoprotected by immersion in 30% (w/v) sucrose in PBS. The brains were then embedded in Cryomatrix embedding resin (Shandon, cat. #6769006), snap frozen in liquid nitrogen and stored at −80°C. 40µm coronal brain cryosections were collected at −18°C and stored as free-floating sections in 0.01% w/v sodium azide (Sigma-Aldrich, cat. #S2002) in PBS. Coronal brain sections spanning approximately −2.255mm to -1.355mm Bregma were selected for immunohistochemistry [Allen Brain Atlas; (Lein et al., 2007)]. Fixed retinae were dissected from eye cups and flat-mounted onto microscope slides for imaging.

### Immunohistochemistry

Antibody manufacturers, host species and dilutions are provided in Supplementary Table 2. Free-floating cryosections were incubated in 100% methanol at −20°C for 10 min, washed in PBS (3 x 5 min, anti-MBP sections only), permeabilised in 0.3% Triton-X (Sigma-Aldrich, cat. #X100-500ML) in PBS for 40 min, blocked with DAKO Serum-Free Protein Block (Agilent, cat. #X0909) for 60 min and incubated overnight at 4°C with primary antibodies diluted in DAKO Antibody Diluent (Agilent, cat. #S080983-2). Sections were washed in PBS (3 x 5 min) prior to incubation with secondary antibody (including lectin) diluted in DAKO Antibody Diluent, at room temperature (∼21C°) for 2h. Sections were stained with DAPI (ThermoFisher, cat. #D5371) for 5 mins, washed with PBS (2 x 5 min), treated with TrueBlack® (Biotium, cat. #23007, NG2DsRed brain sections only), and mounted onto DAKO IHC Microscope Slides (Agilent, cat. #K802021-2) with DAKO Fluorescent Mounting Medium (Agilent, cat. #S302380-2).

### Image acquisition

For each mouse, we imaged one coronal brain section per staining combination. Sections were imaged at 20x magnification on an VS120 Slide Scanner (Olympus, Japan) at excitations of 358nm (blue channel), 488nm (green channel), 594nm (red channel) and 647nm (far-red channel). Each section was imaged in the extended focus imaging mode, over a z-stack distance of 15µm with 3µm intervals. Retinas were imaged using a VS200 Virtual Slide System (Olympus, Japan) at 20x magnification, collecting images spanning a 40µm z-depth at 1.18μm intervals, using the extended focal imaging settings. Optimum exposure times were determined manually for each fluorophore and kept consistent for all images across the cohort.

### Quantification of pericyte density and recombination

To determine the proportion (%) of PDGFRβ^+^ pericytes that underwent cre-mediated transgene recombination, images were opened and blinded in QuPath (v0.4.3) (Bankhead et al., 2017) and regions of interest (ROIs) defined using the Allen Brain Atlas (Lein et al., 2007). ROIs were identified in one brain hemisphere. 0.5mm^2^ ROIs were placed within the somatosensory cortex (spanning layers 2/3 to 6) and thalamus. Within the hippocampus, ROIs were manually traced around Cornu Ammonis (CA)2 and CA3. The corpus callosum ROI was traced from the dorsal-ventral midline to the lateral ventricle. Spinal cord ROIs were traced around the entire grey matter (one section per mouse). For retinae, 0.5mm^2^ ROIs were counted (one section per mouse). PDGFRβ^+^ cells were classified as pericytes if they contained a nucleus (DAPI^+^), colocalised with the microvasculature and had a “bump-on-a-log” morphology (Attwell et al., 2016). Any mural cells on large vessels (>10μm) were excluded from analysis. PDGFRβ^+^ pericytes were manually scored as yellow fluorescent protein (YFP)^+^ or YFP-neg cells within each ROI. YFP^+^ / PDGFRβ-neg cells were also counted.

### Analysis of blood vessel characteristics

Blood vessel length, density and width were determined using FIJI-ImageJ. Briefly, 650 x 650µm ROIs were placed over the somatosensory cortex, thalamus, and hypothalamus of coronal brain confocal images showing anti-CD31 labelling. Images were segmented and vessel characteristics determined using a custom-built ImageJ macro as previously described (Morris et al., 2023b).

To quantify smooth muscle cell coverage of large vessels, images were blinded in QuPath and a ROI placed to define the somatosensory cortex in one hemisphere of a coronal brain section per mouse. Within the ROI, vessels labelled with tomato lectin were classified as large vessels if their diameter was ≥10μm (Hall et al., 2014) at their widest point. Each arteriole segment was manually classified as αSMA^+^ or αSMA-neg, and then a pixel classifier used to determine the area of the traced arterioles that expressed PDGFRβ. Basement membrane coverage of the vasculature was determined by labelling with tomato lectin and anti-CD31. As other cell types also bind tomato lectin, such as microglia (Villacampa et al., 2013), vascular-and parenchymal-tomato lectin staining were segregated by applying pixel classifiers that distinguished tomato lectin labelling (basement membrane) coincident with CD31 labelling (vessel only) using artificial neural network machine learning within QuPath. ICAM-1 area was determined by training a random trees machine learning classifier in QuPath, which measured ICAM-1 immunoreactivity area across the entire somatosensory cortex of one hemisphere per mouse.

For vessel lumen area analysis, separate ROIs encompassing the somatosensory cortex, hippocampus, thalamus, or hypothalamus (in both hemispheres) were overlaid onto confocal images of coronal brain sections from FITC-vascular casted mice labelled with anti-CD31. Trained pixel classifiers were created for the anti-CD31 labelling and FITC vascular cast using the random trees and artificial neural network machine learning within QuPath respectively. The anti-CD31 classifier was used to create an object which contained only the vasculature. The vascular cast classifier was then run within this object to detect the vascular area covered by the vessel cast to give a measure of vessel lumen area.

### Quantification of albumin leakage

For assessment of blood-brain barrier integrity, albumin leakage site density was quantified using QuPath. Images were blinded and ROIs were drawn in the somatosensory cortex and thalamus (one ROI per region per mouse) of coronal brain sections labelled with anti-albumin and anti-CD31. Albumin leakage sites were identified and counted by their close colocalization to CD31^+^ vessels and their cloud-like diffusion staining pattern. Any anti-albumin immunoreactivity which was contained within the anti-CD31 labelling was considered as residual albumin contained within the vessel and so not counted.

### Quantification of Hypoxyprobe^+^ cell density

To quantify the number of hypoxic cells, Hypoxyprobe^+^ cells were manually counted in QuPath. Images were blinded and ROIs drawn that encompassed the hippocampi (two ROIs per section; one section per mouse). A low threshold cell detection detected all DAPI^+^ nucleated cells. As the anti-pimonidazole antibody is pre-conjugated with FITC, and pimonidazole-treated mice were FITC vessel cast), cells were classified as Hypoxyprobe^+^ cells when they were labelled with the FITC pre-conjugated antibody and an Alexa Fluor™ 647 anti-mouse secondary (Invitrogen, cat. #A31571).

### Quantification of glial cell traits

We separately measured the brain area covered by microglia, astrocytes or oligodendrocyte progenitor cells (OPCs), following anti-IBA1, anti-GFAP and anti-PDGFRα labelling, respectively. Using QuPath, a ROI was drawn that encompassed the somatosensory cortex, corpus callosum, hippocampus, thalamus, or hypothalamus in a single hemisphere per imaged brain section. A pixel classifier was created using either machine learning (random trees - IBA1, ANN_MLP - PDGFRα) or single threshold (GFAP) classifiers to detect the area covered by fluorescent pixels. Area and density of microglial clustering was determined by detecting anti-IBA1 immunolabelling using a higher threshold pixel classifier to only detect regions of high immunoreactivity (microglial clusters). Once anti-IBA1 area had been detected, IBA1^+^ areas were converted to objects with a size exclusion of 1200µm^2^ (excluding single activated microglia which had an area of ∼200µm^2^). These large IBA1 detections were counted, and area quantified to determine cluster area and density.

### Statistical analyses

All statistical analyses were performed using GraphPad Prism 10.0.3. Normality was assumed for data sets with n ≤ 3 mice. For datasets n > 3 mice, normality was formally assessed by a Shapiro-Wilk test. If datasets were normal, single factor comparisons were analysed with an unpaired t-test. Comparisons of two variables were analysed with a two-way ANOVA with Tukey’s or Šidák’s multiple comparisons test. Non-parametric datasets were analysed with a Mann-Whitney test. Statistical outcomes including overall p-values are provided in the corresponding figure legends. Data are presented as mean ± standard deviation unless otherwise stated. *p* < 0.05 was considered statistically significant.

## Results

### Tamoxifen administration induces pericyte YFP expression in transgenic mice

To determine the extent and specificity of transgene recombination achieved using *Pdgfrβ-CreER^T2^* mice, we crossed these mice with the cre-sensitive *Rosa26-YFP* reporter strain to produce *Pdgfrβ-CreER^T2^ :: Rosa26-YFP* (βCre-YFP) mice (Figure 1A). βCre-YFP mice received: (i) one 100mg/kg dose; (ii) one 300mg/kg dose, or (iii) four consecutive daily doses of 300mg/kg tamoxifen (Tx) to activate cre-recombinase. 7-days later (Tx+7), brain tissue was collected and used to generate coronal brain cryosections in which yellow fluorescent protein (YFP) was detected in pericytes expressing PDGFRβ (Figure 1B).

**Figure 1.**
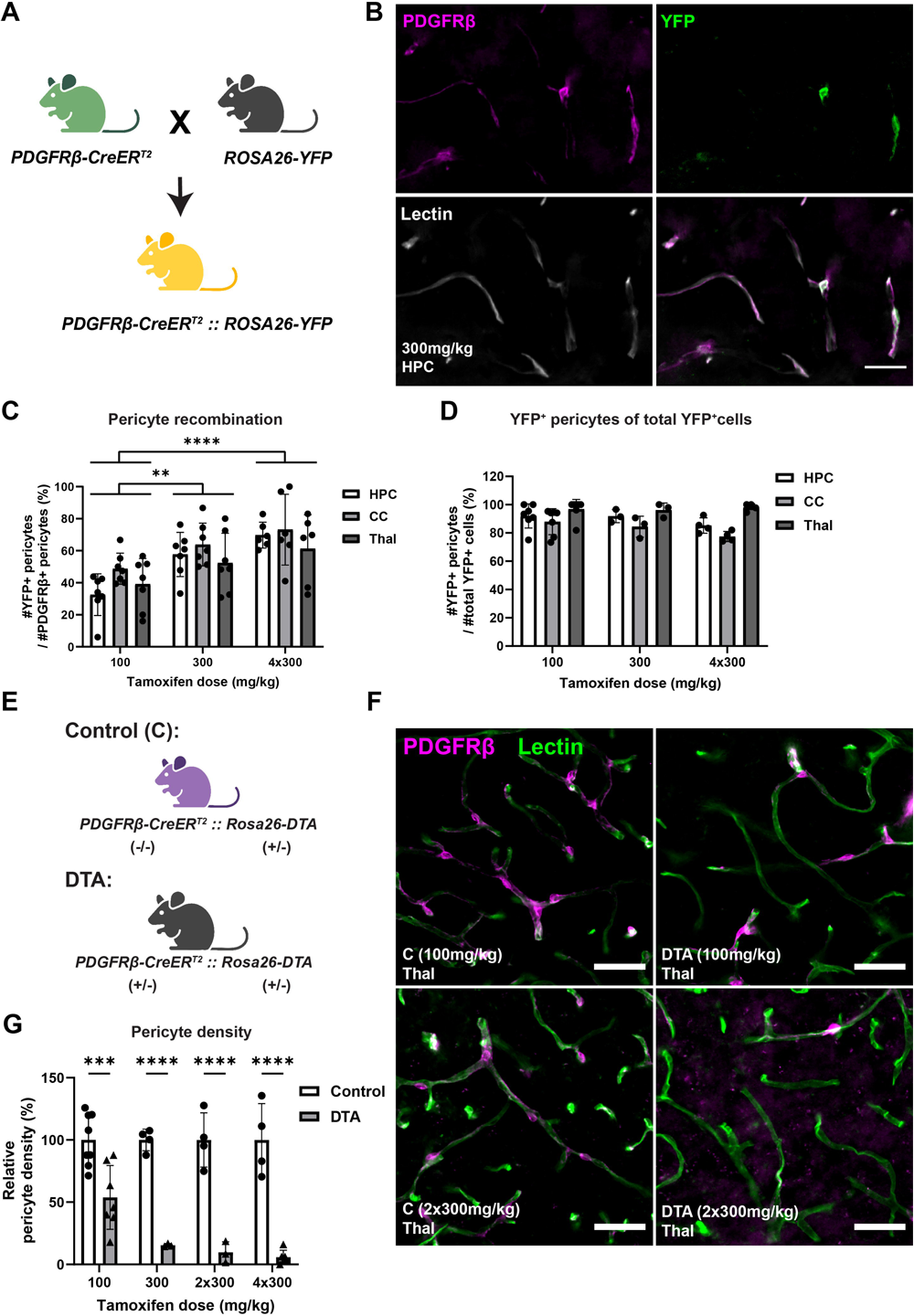
YFP expression and pericyte ablation in the brains of transgenic mice 7-days following tamoxifen administration. **A)** Breeding strategy for producing *PDGFRβCreER^T2^ :: Rosa26YFP* transgenic mice. **B)** Immunofluorescent image of tomato lectin (white), yellow fluorescent protein (YFP, green) and platelet-derived growth factor receptor beta (PDGFRβ, magenta) labelling of pericytes on blood vessels in *PDGFRβCreER^T2^ :: Rosa26YFP* mice. Scale = 25µm. **C)** Percentage of recombined (YFP^+^) pericytes out of all PDGFRβ^+^ pericytes in different brain regions. HPC = hippocampus, CC = corpus callosum, Thal = thalamus. Two-way ANOVA: tamoxifen dose: F (2, 51) = 16.17, *p* < 0.0001; brain region: F (2, 51) = 2.718, *p* =0.0755; interaction: F (4, 51) = 0.5045, *p* = 0.7325). **D)** Percent YFP^+^ pericytes of total YFP^+^cells. **E)** Control and DTA *PDGFRβCreER^T2^ :: Rosa26DTA* mice genotypes. **F)** Immunofluorescent images of thalamic brain sections labelled with tomato lectin (basement membrane, green) and PDGFRβ (mural cells including pericytes, magenta) from *PDGFRβCreER^T2^ :: Rosa26DTA* mice without (control) and with (DTA) the Cre transgene 7-days post tamoxifen administration. Scale = 40µm. **G)** Relative pericyte density quantification in the thalamus of control and DTA mice after various doses of tamoxifen. Two-way ANOVA: tamoxifen dose: F (3, 31) = 4.035, *p* = 0.0156; genotype: F (1, 31) = 144.3, *p* < 0.0001; interaction: F (3, 31) = 0.4.305, *p* = 0.0156. Data shown as mean ± SD, with individual data points for each animal. Two-way ANOVA with Tukey’s or Šidák’s multiple comparisons test. ** *p* < 0.01, *** *p* < 0.001, **** *p* < 0.0001. Panels A and E were created with BioRender.

The proportion of PDGFRβ^+^ pericytes that became YFP-labelled was partially dependent on Tx dose. After receiving a single dose of 100mg/kg Tx, YFP labelling was present in ∼33% of hippocampal, ∼49% of callosal and ∼39% of thalamic pericytes (Figure 1C). Cre-mediated recombination was significantly increased when βCre-YFP mice received a single dose of 300mg/kg Tx, as ∼58% of pericytes in the hippocampus, ∼64% in the corpus callosum and ∼52% in the thalamus were YFP-labelled (*p* = 0.0015). Four consecutive daily doses of 300mg/kg Tx did not induce a further increase in recombination in any of the brain regions analysed, when compared to a single dose of 300mg/kg Tx. Irrespective of the Tx dose, the vast majority of YFP^+^ cells (>90%) in the hippocampus, corpus collosum, and thalamus were pericytes (Figure 1D). While a small subset of YFP^+^ parenchymal cells expressed GFAP and had an astrocytic morphology (Supplementary Figure 1), these data confirm that *Pdgfrβ- CreER^T2^* mice can be used to reliably target pericytes in the brain, and that Tx dose can influence the proportion of the pericyte population affected.

### Tamoxifen administration induces pericyte ablation in transgenic mice

We used the knowledge gained from analysing βCre-YFP mice to titrate the genetic ablation of pericytes. *Pdgfrβ-CreER^T2^* mice were crossed with *Rosa26-DTA* mice to generate *Pdgfrβ- CreER^T2^ :: Rosa26-DTA* (βCre-DTA) mice (Figure 1E). Control (*cre* -/-, *Rosa26-DTA* +/-) and DTA (*cre* +/-, *Rosa26-DTA* +/-) mice received: (i) one 100mg/kg dose (DTA^100mg/kg^); (ii) one 300mg/kg dose (DTA^300mg/kg^), (iii) two consecutive daily doses of 300mg/kg (DTA^2×300mg/kg^) or (iv) four consecutive daily doses of 300mg/kg (DTA^4×300mg/kg^) Tx and brain tissue was collected at Tx+7. PDGFRβ^+^ pericytes were detected in all brain regions examined in coronal cryosections from control mice (Supplementary Figure 2). The density of PDGFRβ^+^ pericytes appeared visually decreased in the thalamus of DTA mice (Figure 1F). Quantification revealed that relative to control mice, thalamic pericyte density was reduced by ∼46% (*p* = 0.0002) in DTA^100mg/kg^ mice, ∼85% in DTA^300mg/kg^ (*p* < 0.0001), ∼90% in DTA^2×300mg/kg^ (*p* < 0.0001) and ∼94% in DTA^4×300mg/kg^ (*p* < 0.0001) mice (Figure 1G). Pericytes were also ablated in other grey matter regions (Figure 2A), with pericyte density being mildly reduced in the somatosensory cortex (∼44%, *p* < 0.0001), hippocampus (∼58%, *p* < 0.0001), and hypothalamus (∼44%, *p* = 0.0056) of DTA^100mg/kg^ mice (Figure 2B). Pericyte density was not significantly reduced in the corpus callosum of DTA^100mg/kg^ mice compared to controls (Figures 2A,B). In DTA^2×300mg/kg^ mice, pericyte density was not only more strongly reduced in each of the grey matter regions examined (≥ 80%) but was also significantly reduced in the corpus callosum (∼80%, *p* = 0.0002; Figure 2C).

**Figure 2.**
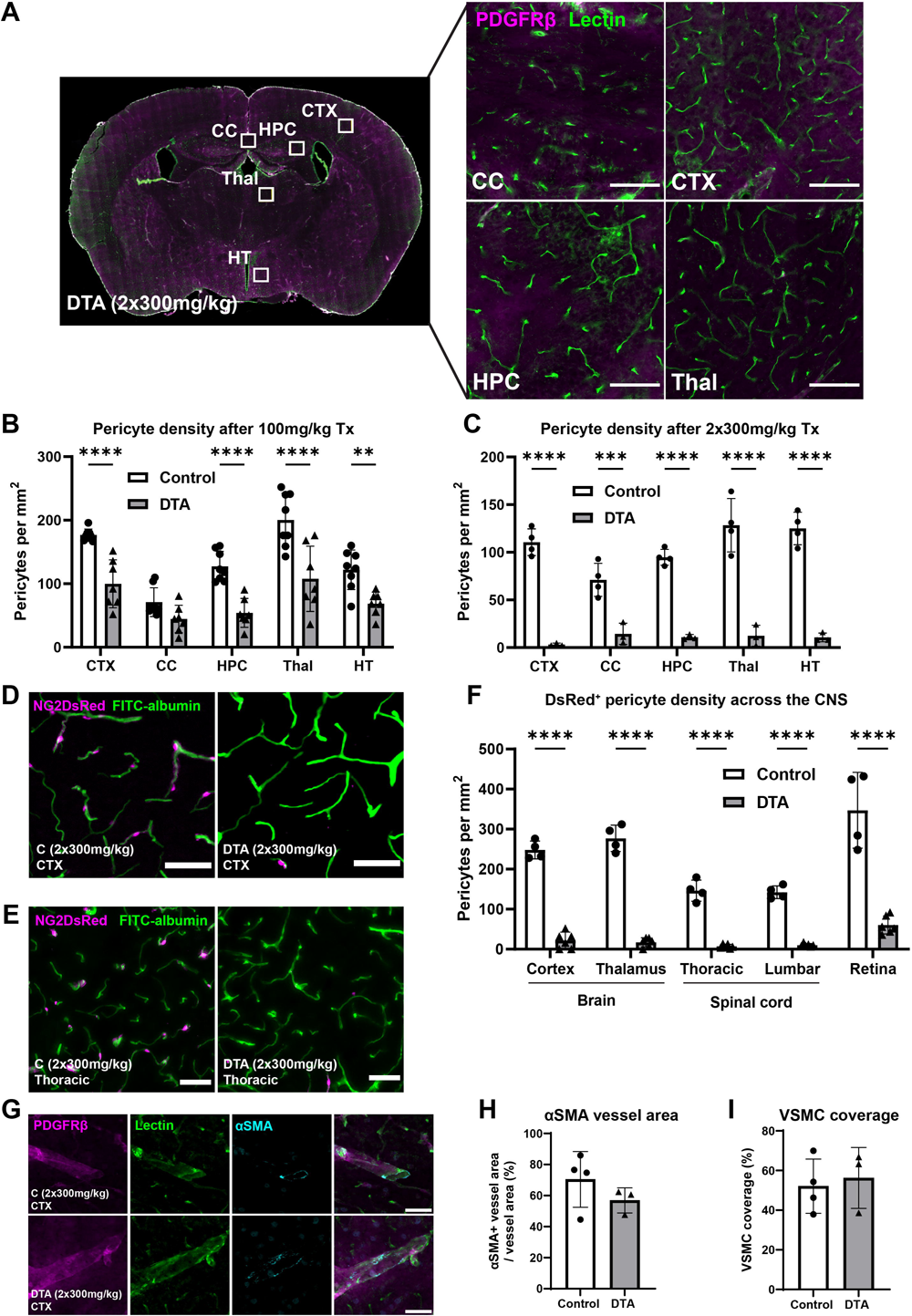
Pericyte ablation occurs across the central nervous system in transgenic mice 7-days following tamoxifen administration. **A)** PDGFRβ (mural cells including pericytes) and tomato lectin (basement membrane) immunolabelling in the somatosensory cortex (CTX), corpus callosum (CC), hippocampus (HPC) and thalamus (Thal) of a *PDGFRβCreER^T2^ :: Rosa26DTA* mouse after DTA activation. Scale = 100µm. **B-C)** Pericyte density across different brain regions in control and DTA mice after B) 100mg/kg (two-way ANOVA: brain region: F (4, 65) = 24.55, *p* < 0.0001; genotype: F (1, 65) = 84.87, *p* < 0.0001; interaction: F (4, 65) = 2.637, *p* = 0.0418) or C) 2×300mg/kg (two-way ANOVA: brain region: F (4, 25) = 4.000, *p* = 0.0121; genotype: F (1, 25) = 358.3, *p* < 0.0001; interaction: F (4, 25) = 5.013, *p* = 0.0042) tamoxifen (Tx). HT = hypothalamus. **D)** Fluorescent images of NG2DsRed (magenta) and FITC-albumin (green) from the somatosensory cortex of *PDGFRβCreER^T2^ :: Rosa26DTA :: NG2DsRed* mice. Scale = 50µm. **E)** Fluorescent images of NG2DsRed (magenta) and FITC-albumin (green) from the thoracic spinal cord of control and DTA mice. Scale = 50µm. **F)** Quantification of pericyte density from NG2-DsRed fluorescence across the CNS of control and DTA mice. Two-way ANOVA: CNS region: F (4, 40) = 27.56m *p* < 0.0001; genotype: F (1, 40) = 522.7, *p* < 0.0001; interaction: F (4, 40) = 11.78, *p* < 0.0001. **G)** Immunofluorescent images of vascular smooth muscle cells labelled with alpha smooth muscle actin (αSMA, cyan) and platelet-derived growth factor beta (PDGFRβ, magenta), and tomato lectin (green) to label the vasculature in control and DTA mice. Scale = 40µm. **H-I)** Quantification of J) αSMA^+^ large vessel area and K) vascular smooth muscle cell (VSMC) coverage in control and DTA mice. Data shown as mean ± SD, with individual data points for each animal. Two-way ANOVA with Šidák’s multiple comparisons test and unpaired t tests. ** *p* < 0.01, *** *p* < 0.001, **** *p* < 0.0001.

To confirm that these data reflected pericytes loss, rather than the downregulation of PDGFRβ protein, βCre-DTA mice were crossed with NG2-DsRed mice, to fluorescently tag pericytes with the DsRed protein. As NG2 (neuron-glial antigen 2) is also expressed in OPCs, we initially quantified the relative DsRed labelling of pericytes and OPCs in NG2-DsRed mice. Consistent with our previous report (Morris et al., 2023a), only ∼7% of OPCs were labelled with DsRed across all brain regions analysed (Supplementary Figure 3A-C). OPCs that were DsRed^+^ also had a significantly less intense nuclear DsRed signal compared to pericytes (Supplementary Figure 3D; *p* < 0.0001). *Pdgfrβ-CreER^T2^ :: Rosa26-DTA :: NG2DsRed* [DTA-DsRed (*cre* +/-, *Rosa26-DTA* +/-, *NG2DsRed* +/-), control (*cre* -/-, *Rosa26-DTA* +/-, *NG2DsRed* +/-)] mice received 2 consecutive daily 300mg/kg doses of Tx. When the number of DsRed^+^ pericytes was quantified at Tx+7, DsRed^+^ pericyte density was reduced by ∼91% (*p* < 0.0001) in the somatosensory cortex and ∼97% (*p* < 0.0001) in the thalamus of DTA-DsRed^2×300mg/kg^ mice, compared to controls (Figure 2D,F). Other studies have previously shown that brain and retinal pericytes are ablated in inducible DTA (diphtheria toxin fragment A) / DTR (diphtheria toxin receptor) pericyte depletion models (Eilken et al., 2017, Nikolakopoulou et al., 2019), but spinal cord pericyte ablation has not been explored. We determined that pericyte density was reduced by ∼95% in the thoracic and ∼93% in the lumbar spinal cord of DTA-DsRed^2×300mg/kg^ mice compared to controls (Figure 2E,F; both *p* < 0.0001). Pericyte density was also reduced by ∼83% in the retina of DTA-DsRed^2×300mg/kg^ mice compared to controls (Supplementary Figure 4, Figure 2F, *p* < 0.0001;).

As vascular smooth muscle cells (VSMCs) also express PDGFRβ (Nikolakopoulou et al., 2019), we explored the possibility that VSMCs were also ablated in Tx+7 DTA mice. Coronal brain sections from control and DTA mice were processed to detect PDGFRβ, tomato lectin (labelling the basement membrane of blood vessels) and alpha smooth muscle actin (αSMA) (Figure 2G). αSMA is highly expressed by VSMCs and expressed at a low level in pericytes *in vivo* (Smyth et al., 2018, Vanlandewijck et al., 2018). The proportion of tomato lectin^+^ large vessels (arterioles and venules, ≥10μm in diameter) labelled with αSMA^+^ was equivalent in control and DTA^2×300mg/kg^ mice (Figure 2H). Large vessel coverage of VSMCs was also unchanged (Figure 2I). Together, these data suggest that ablation was restricted to pericytes and not VSMCs in Tx+7 DTA mice.

### Pericyte ablation reduces activity and causes a rapid-onset motor impairment in mice

Having identified Tx doses that produce a moderate (100mg/kg) or severe (≥300mg/kg) level of pericyte ablation, we next explored the impact of varying levels of pericyte ablation on mouse behaviour. Each Tx+7 control, DTA^100mg/kg^, DTA^300mg/kg^, or DTA^2×300mg/kg^ mouse was placed in an open field arena for 10 min and their voluntary movement was recorded (Figure 3A). Quantification of the total distance travelled across the 10 min period revealed that DTA^100mg/kg^ and DTA^300mg/kg^ and travelled ∼45% (*p* = 0.0010) and ∼51% (*p* = 0.0030) less than control mice, whereas DTA^2×300mg/kg^ travelled ∼85% (*p* < 0.0001) less than controls (Figure 3B). This phenotype can be partially explained by a reduction in the time that DTA mice spent moving around the arena (movement time; Figure 3C), a decrease in their maximum acceleration (Supplementary Figure 5A), and an increase in the amount of time spent inactive (Figure 3C, Supplementary Figure 5B). By categorising movement into activity states, we determined that DTA^100mg/kg^ mice spent ∼53% less time in a moderate activity state and ∼23% more time in a low activity state compared to controls (both *p* < 0.0001; Figure 3D). This was exacerbated in DTA^2×300mg/kg^ mice, as they spent ∼85% less time spent in a moderate activity state and ∼48% more time in a low activity state than control mice (both *p* < 0.0001; Figure 3E). Severe pericyte ablation was also associated with a reduction in body weight and poorer health scores (Supplementary Figure 6). Collectively, these data suggest that pericyte ablation induces hypoactivity.

**Figure 3.**
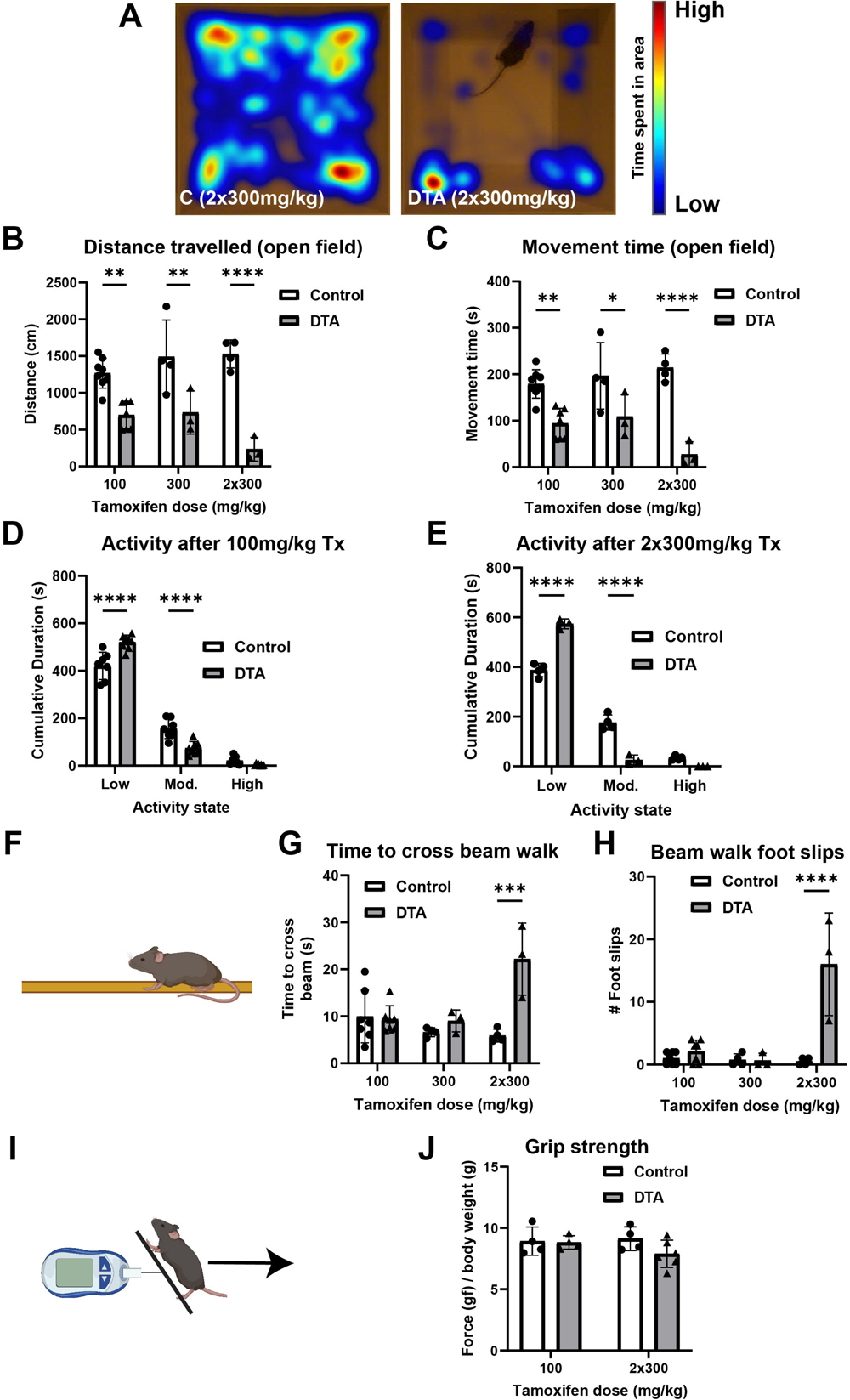
Mice with reduced pericytes exhibit fatigue and motor impairment 7-days after tamoxifen administration. **A)** Heatmaps of the relative time spent across the open field arena over 10 mins. **B-E)** Quantification of B) distance travelled (two-way ANOVA: tamoxifen dose: F (2, 23) = 1.345, *p* = 0.2802; genotype: F (1, 23) = 69.47, *p* < 0.0001; interaction: F (2, 23) = 4.492, *p* = 0.0226), C) time spent moving (two-way ANOVA: tamoxifen dose: F (2, 23) = 1.086, *p* = 0.3543; genotype: F (1, 23) = 55.93, *p* < 0.0001; interaction: F (2, 23) = 4.125, *p* =0.0295) and D-E) activity of control and DTA mice after administration with varying doses of tamoxifen [D, 100mg/kg (two-way ANOVA: activity state: F (2, 39) = 766.1, *p* < 0.0001; genotype: F (1, 39) = 0.0005148. *p* = 0.9820; interaction: F (2, 39) = 28.26, *p* < 0.0001); E, 2×300mg/kg (two-way ANOVA: activity state: F (2, 15) = 932.6, *p* < 0.0001; genotype: F (1, 15) = 0.002876, *p* = 0.9579; interaction: F (2, 15) = 111.4, *p* < 0.0001)] in a 10-minute open field task. Mod. = moderate. Tx = tamoxifen. **F)** Visual representation of the beam walk task. **G)** Time to cross beam (two-way ANOVA: tamoxifen dose: F (2, 22) = 4.213, *p* = 0.0282; genotype: F (1, 22) = 13.30, *p* = 0.0014; interaction: F (2, 22) = 9.877, *p* = 0.0009) and **H)** number of slips (two-way ANOVA: tamoxifen dose: F (2, 22) = 16.88, *p* < 0.0001; genotype: F (1, 22) = 25.13, *p* < 0.0001; interaction: F (2, 22) = 18.82, *p* < 0.0001) by control and DTA mice challenged with the beam walk task. **I)** Visual representation of the grip strength task. **J)** Grip strength relative to body weight in control and DTA mice. Data shown as mean ± SD, with individual data points for each animal. Two-way ANOVA with Šídák’s multiple comparisons test and unpaired t-tests. * *p* < 0.05, ** *p* < 0.01, *** *p* < 0.001, **** *p* < 0.0001. Panels F and I were created with BioRender.

After observing that DTA mice were less active, we aimed to determine whether they also showed signs of impaired motor coordination. In the beam walk task (Figure 3F), control, DTA^100mg/kg^ and DTA^300mg/kg^ mice took similar amounts of time to cross the beam (Figure 3G). However, DTA^2×300mg/kg^ mice took on average ∼16 seconds longer to cross the beam than controls (Figure 3G; *p* = 0.0001). Further, an assessment of foot slips showed that DTA^2×300mg/kg^ mice slipped ∼16 times while control mice rarely (<1) made foot slip errors (*p* < 0.0001). By contrast, the performance of DTA^100mg/kg^ and DTA^300mg/kg^ mice was not different to that of controls (Figure 3H). Total steps taken was equivalent across all groups (Supplementary Figure 5C). These data demonstrate that ablation of >80% of pericytes leads to a loss of motor coordination. By performing a grip strength assessment to measure muscular strength (Figure 3I), we determined that relative muscle strength was equivalent in control, DTA^100mg/kg^ and DTA^2×300mg/kg^ mice (Figure 3J), suggesting that muscular weakness was not the underlying cause of hypoactivity or impaired motor coordination following pericyte ablation.

### Pericyte ablation does not change vessel structure, but alters vessel lumen size and blood-brain barrier leakage

After detecting significant reductions in activity and motor function in DTA mice, we decided to explore potential pathological mechanisms which might be driving this behavioural phenotype. We first investigated vascular integrity, as pericytes play an integral role in maintaining and regulating vessel function in the brain (Brown et al., 2019). To determine whether vessel structure was altered by pericyte ablation, coronal brain sections from Tx+7 control and DTA^2×300mg/kg^ mice were labelled with the endothelial cell marker CD31 (cluster of differentiation 31, Figure 4A). Mean vessel width (Figure 4B), vessel density (Figure 4C) and vessel length (Figure 4D) were equivalent between treatment groups in all brain regions analysed. Pericytes also contribute to the maintenance and remodelling of other vascular components such as the basement membrane (Wang et al., 2012). To determine whether pericyte ablation changes basement membrane integrity, brain sections were labelled with tomato lectin to label glycoproteins in the basement membrane (Figure 4A) (Reitsma et al., 2007). Basement membrane coverage was unchanged in the brains of DTA^2×300mg/kg^ mice compared to controls (Figure 4E, Supplementary Figure 7). As pericytes regulate neuroinflammation in the brain by controlling the expression of leukocyte adhesion molecules (LAMs) on the endothelium (Torok et al., 2021), we next labelled brain sections to detect tomato lectin and ICAM-1 (Figure 4F). ICAM-1 area was increased by ∼107% (*p* = 0.0039) in DTA^2×300mg/kg^ mice compared to controls (Figure 4G), suggestive of an increase in neuroinflammation.

**Figure 4.**
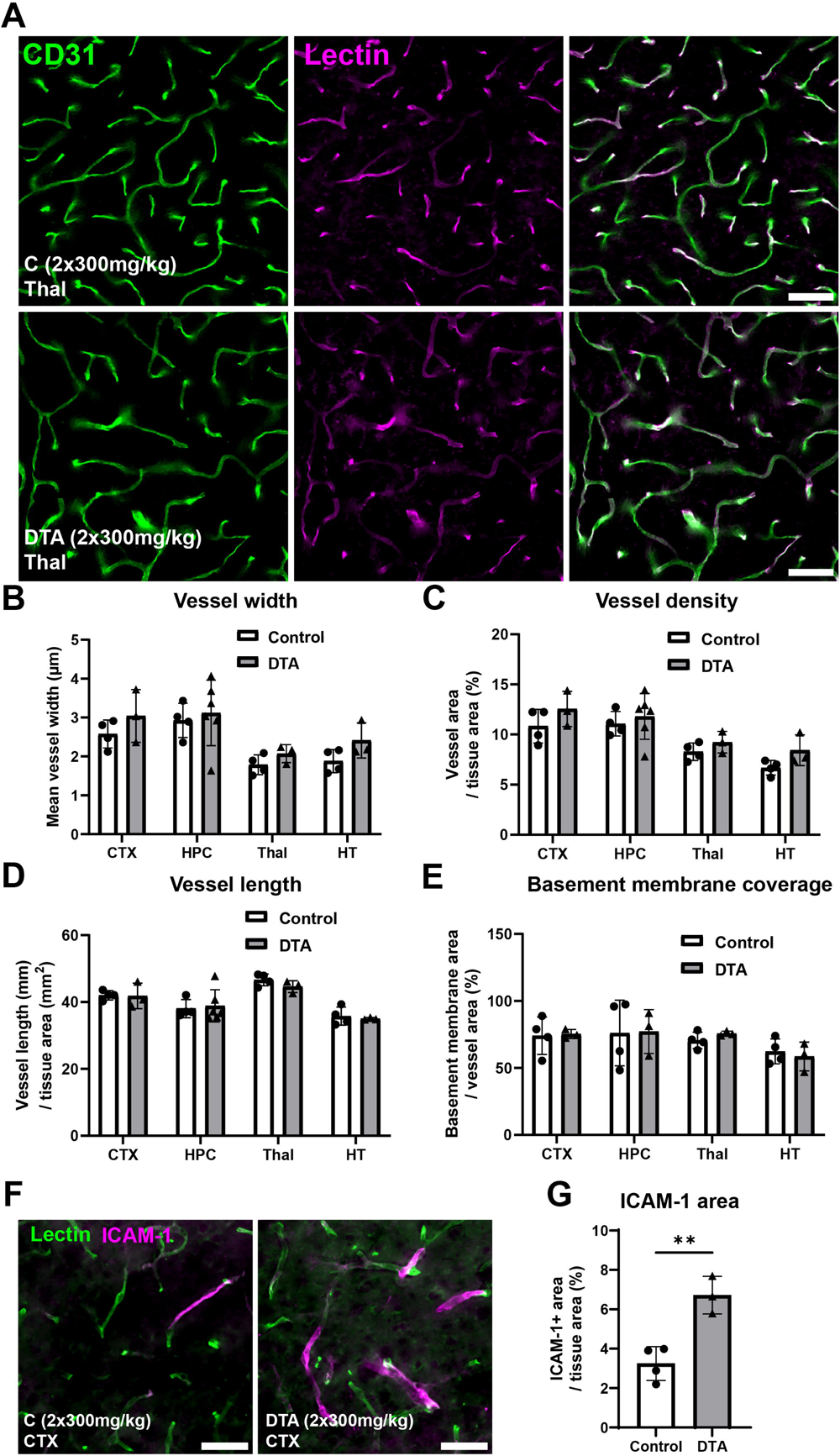
Brain vessel density and basement membrane coverage remains unchanged while ICAM-1 expression increases 7-days post-pericyte ablation. **A)** Immunofluorescent images of CD31 (cluster of differentiation 31, green) and tomato lectin (magenta) labelling in the thalamus of control and DTA mice. Scale = 50µm. **B-E)** Vascular structure indicated by B) mean vessel width, C) vessel density, D) vessel length and E) basement membrane coverage in the somatosensory cortex (CTX), hippocampus (HPC), thalamus (Thal) and hypothalamus (HT) of control and DTA mice. **F)** Immunofluorescent images of tomato lectin (green) and ICAM-1 (intercellular adhesion molecule 1, magenta) in control and DTA mice. **G)** ICAM-1 area in the somatosensory cortex of control and DTA mice. Unpaired t-test: t = 5.059, df = 5, *p* = 0.0039. Data shown as mean ± SD, with individual data points for each animal. Unpaired t test. ** *p* < 0.01.

Pericytes regulate blood flow by contracting and dilating capillaries to modulate vessel width (Hall et al., 2014, Dessalles et al., 2021). We hypothesised that a loss of pericytes may lead to reduced tone on capillaries, resulting in changes in luminal diameter and area. To determine if luminal vessel diameter was changed by pericyte ablation, βCre-DTA mice were perfused with a FITC-albumin gel to cast the vessel lumen (Figure 5A) and cryosections stained to detect the surrounding CD31^+^ endothelial cell layer (Supplementary Figure 8). Irrespective of the brain region analysed, we consistently found that vessel lumen area was increased in DTA^2×300mg/kg^ mice compared to controls (two-way ANOVA: genotype F(1,35) = 18.47, *p* = 0.001), however, this was most striking in the hippocampus (*p* = 0.0112, Figure 5B). In DTA^2×300mg/kg^ mice, mean lumen width was increased by ∼39% (*p* = 0.0457) and ∼35% (*p* = 0.0490) in the hippocampus and thalamus, respectively (Figure 5C). These data suggest that pericyte loss reduces capillary tone in some brain regions.

**Figure 5.**
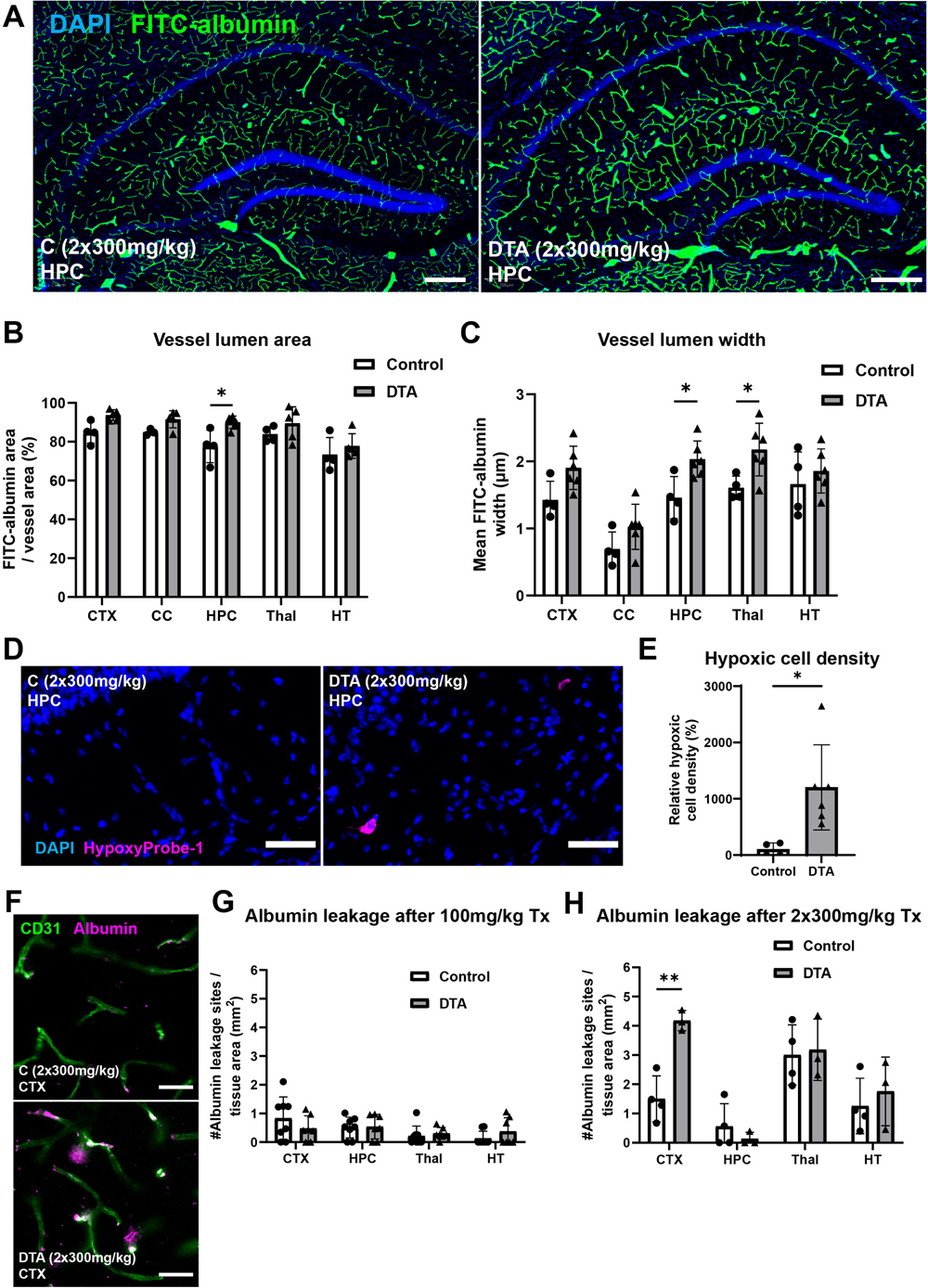
Vessel lumen diameter, hypoxic cells and BBB leakage is increased 7-days post-pericyte ablation. **A)** Immunofluorescent images of cell nuclei (DAPI, blue) and vessel lumen (FITC-albumin gel casting, green) in the hippocampus (HPC) of control and DTA mice. Scale = 250µm. **B-C)** Quantification of B) vessel lumen area (two-way ANOVA: brain region: F (4, 35) = 7.784, *p* = 0.0001; genotype: F (1, 35) = 18.47, *p* = 0.0001; interaction: F (4, 35) = 0.6787, *p* = 0.6114) and C) vessel lumen width (two-way ANOVA: brain region: F (4, 40) = 15.37, *p* < 0.0001; genotype: F (1, 40) = 20.77, *p* < 0.0001; interaction: F (4, 40) = 0.6065, *p* = 0.6603) of control and DTA mice in the somatosensory cortex (CTX), corpus callosum (CC), hippocampus (HPC), thalamus (Thal) and hypothalamus (HT). **D)** Immunofluorescent labelling of DAPI (nuclei, blue) and Hypoxyprobe-1 (magenta) in the HPC of control and DTA mice. Scale = 50µm. **E)** Quantification of Hypoxyprobe-1 positive cells in the HPC of control and DTA mice. Mann-Whitney test, *p* = 0.0095. **F)** Immunofluorescent labelling of endothelial cells (CD31, green) and albumin (magenta) in the CTX of control and DTA mice. Scale = 30µm. **G-H)** Quantification of albumin leakage sites in the CTX, Thal, HPC and HT of control and DTA mice [G, 100mg/kg; H, 2×300mg/kg (two-way ANOVA: brain region: F (3, 20) = 14.99, *p* < 0.0001; genotype: F (1, 20) = 4.957, *p* = 0.0376; interaction: F (3, 20) = 4.214, *p* = 0.0183)]. Tx = tamoxifen. Data shown as mean ± SD, with individual data points for each animal. Two-way ANOVA with Šídák’s multiple comparisons. * *p* < 0.05, ** *p* < 0.01.

As pericytes are crucial for regulating and distributing blood flow, we aimed to determine whether pericyte ablation was associated with an increase in brain hypoxia. Hypoxic cells were detected using Hypoxyprobe-1 (HP-1, Figure 5D). DTA^2×300mg/kg^ mice had ∼1086% (*p* = 0.0220) more HP-1^+^ cells compared to controls (Figure 5E). The HP-1^+^ cells expressed somatostatin, identifying them as interneurons (Supplementary Figure 9A), which is consistent with the hypothesis that HP-1+ cells in the brain are interneurons (Nortley et al., 2019). Despite an increase in the number of hypoxic interneurons, this was insufficient to drive a change in the number of somatostatin^+^ cells detected in hippocampus after pericyte ablation (Supplementary Figure 9B-C). As pericytes contribute to the regulation and maintenance of the BBB (Armulik et al., 2010, Daneman et al., 2010), coronal brain sections from βCre-DTA mice were labelled with albumin and CD31 to detect sites of albumin leakage into the brain parenchyma (Figure 5F). Pericyte ablation in the DTA^100mg/kg^ mice did not change overall albumin labelling in the brain, relative to controls (Figure 5G). However, there was an increase of ∼178% in the density of cortical locations with albumin leakage (Figure 5H; *p* = 0.0024). These data suggest pericyte ablation can trigger hypoxia in interneurons and causes BBB leakage in some brain regions.

### Pericyte ablation induces microgliosis and astrogliosis

As pericytes are closely associated with glial populations within the neurovascular unit (Cashion et al., 2023, Morris et al., 2023a), we examined the possibility that pericyte ablation could impact the state of microglia or astrocytes. To investigate the microglial response to pericyte loss, βCre-DTA coronal brain sections were labelled to detect the microglial protein, IBA1 (ionised calcium binding adaptor molecule 1; Figure 6A,B). IBA1^+^ area was equivalent in control and DTA^100mg/kg^ mice (Figure 6C), but significantly elevated in the somatosensory cortex (∼76%, *p* < 0.0001), corpus callosum (∼129%, *p* = 0.0004) and hippocampus (∼100%, *p* < 0.0001) of DTA^2×300mg/kg^ mice (Figure 6D). Microglia also formed dense IBA1^+^ clusters in DTA^2×300mg/kg^ mice (Figure 6B), suggestive of active microgliosis. Quantification of microglial cluster area and cluster density in DTA^2×300mg/kg^ mice revealed that these measures were significantly increased in the somatosensory cortex, hippocampus and hypothalamus, compared to controls (Figure 6E,F). These data suggest that pericyte ablation induces significant microgliosis.

**Figure 6.**
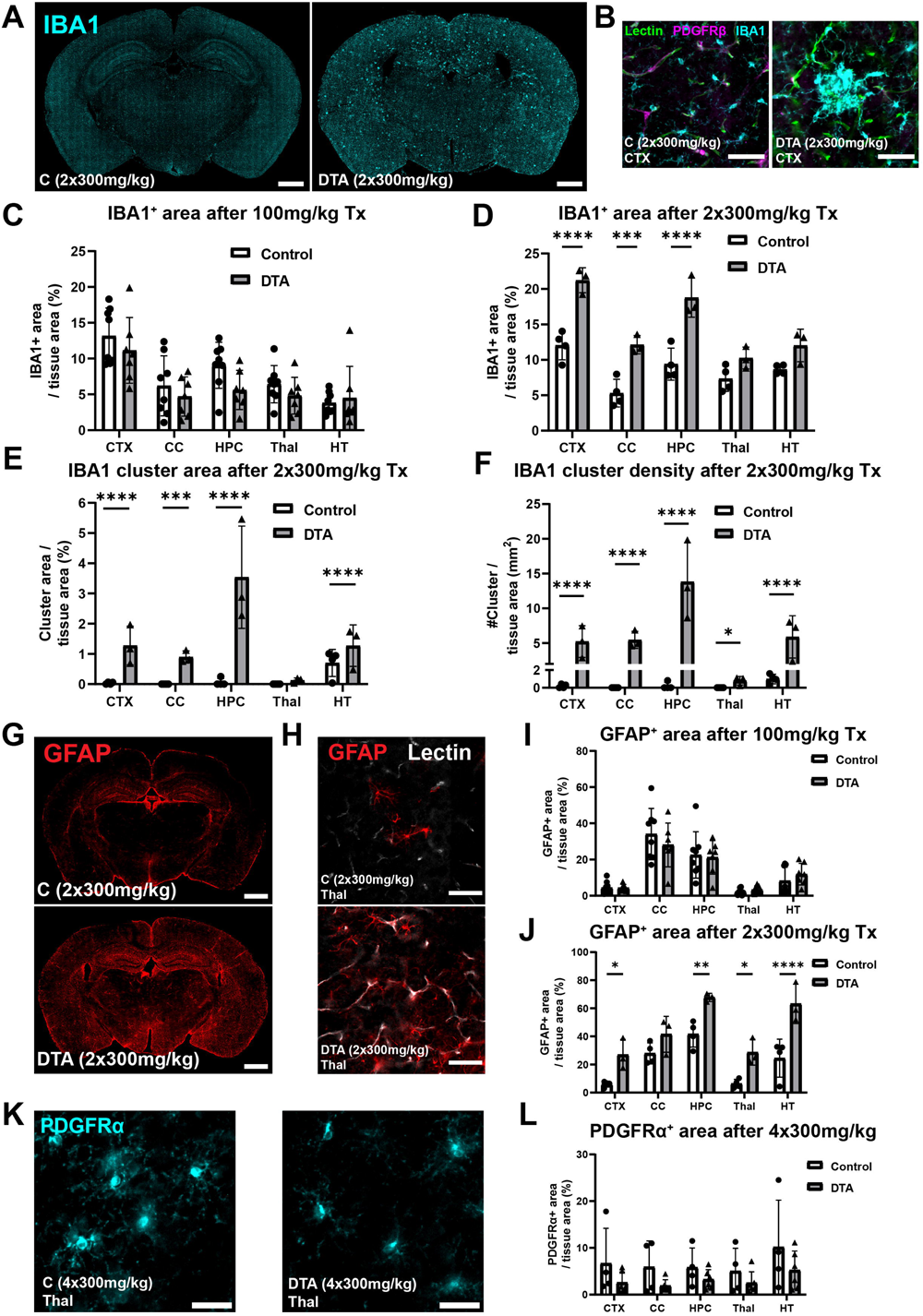
Microgliosis and astrogliosis occur 7-days post-pericyte ablation. **A)** Full coronal brain sections labelled with IBA1 (ionized calcium binding adaptor molecule 1, cyan) representing microglia from control and DTA mice. Scale = 1mm. **B)** Immunofluorescent images of tomato lectin (basement membrane, green), platelet-derived growth factor receptor (PDGFRβ, magenta) and IBA1 (cyan) labelling in the somatosensory cortex of control and DTA mice. Scale = 50µm. **C-F)** Quantification of C-D) IBA1 area [two-way ANOVA: 2×300mg/kg (brain region: F (4, 25) = 23.40, *p* < 0.0001; genotype: F (1, 25) = 95.40, *p* < 0.0001; interaction: F (4, 25) = 4.528, *p* = 0.0069)], E) IBA1 cluster area (two-way ANOVA: brain region: F (4, 25) = 13.13, *p* < 0.0001; genotype: F (1, 25) = 89.37, *p* < 0.0001; interaction: F (4, 25) = 10.86, *p* < 0.0001) and F) IBA1 cluster density (two-way ANOVA: brain region: F (4, 25) = 15.44, *p* < 0.0001; genotype: F (1, 25) = 246.8, *p* < 0.0001; interaction: F (4, 25) = 10.99, *p* < 0.0001) across various brain regions in control and DTA mice (C, 100mg/kg; D-F, 2×300mg/kg). CTX = somatosensory cortex, CC = corpus callosum, HPC = hippocampus, Thal = thalamus, HT = hypothalamus. **G)** Full coronal brain sections labelled with GFAP (glial fibrillary acidic protein, red) representing astrocytes from control and DTA mice. Scale = 1mm. **H)** Immunofluorescent images of thalamic tomato lectin (basement membrane, white) and GFAP (red) labelling in control and DTA mice. Scale = 50µm. **I-J)** Quantification of GFAP area across various brain regions in control and DTA mice [I, 100mg/kg; J, 2×300mg/kg (two-way ANOVA: brain region: F (4, 25) = 21.91, *p* <0.0001; genotype: F (1, 25) = 58.82, *p* < 0.0001; interaction: F (4, 25) = 1.691, *p* = 0.1836)]. **K)** Immunofluorescent labelling of oligodendrocyte progenitor cells (OPCs) with PDGFRα (cyan) in the Thal of control and DTA mice. Scale = 30µm. **L)** Quantification of PDGFRα immunoreactivity (two-way ANOVA: brain region: F (4, 40) = 1.151, *p* = 0.3468; genotype: F (1, 40) = 7.334, *p* = 0.0099; interaction: F (4, 40) = 0.1239, *p* = 0.9730) across various brain regions in control and DTA mice. Tx = tamoxifen. Data shown as mean ± SD, with individual data points for each animal. Log(x+1) transformation (E and F only). Two-way ANOVA with Šídák’s multiple comparisons test. * *p* < 0.05, ** *p* < 0.01, *** *p* < 0.001, **** *p* < 0.0001.

As pericytes also associate with astrocytes in the neurovascular unit (Mishra et al., 2016), coronal brain sections from βCre-DTA were labelled with GFAP (glial fibrillary acidic protein) to label fibrous and / or reactive astrocytes (Figure 6G,H). GFAP immunoreactivity area appeared equivalent in control and DTA^100mg/kg^ mice (Figure 6I), but was significantly elevated in DTA^2×300mg/kg^ mice, in the somatosensory cortex (∼448%, *p* = 0.0335), hippocampus (∼63%, *p* = 0.0056), thalamus (∼452%, *p* = 0.0206), and hypothalamus (∼258%, *p* < 0.0001) compared to controls (Figure 6J).

Cells of the oligodendrocyte lineage are particularly sensitive to the toxic effects of blood proteins (Petersen et al., 2017) and to hypoxia (Káradóttir et al., 2008) and a subset of OPCs closely associate with the microvasculature in the brain and may interact with pericytes (Maki et al., 2015). To determine how OPCs respond to pericyte ablation, coronal brain sections from control and DTA mice were processed to detect platelet-derived growth factor receptor alpha (PDGFRα; Figure 6K), which is highly expressed by OPCs (Rivers et al., 2008). The area of PDGFRα immunoreactivity was significantly lower in DTA^4×300mg/kg^ mice relative to controls, across multiple brain regions (two-way ANOVA: genotype: F (1, 40) = 7.334, *p* = 0.0099) (Figure 6L). However, by staining brain sections to detect myelin basic protein (MBP), we found that pericyte ablation was not associated with an overt change in myelination (Supplementary Figure 10). These data suggest that pericyte ablation is associated with a shift in the microglial and astrocytic state, potentially towards a more neuroinflammatory phenotype. The reduction in PDGFRα immunoreactive area may reflect a reduction in OPC arborisation, which could similarly reflect a shift towards a more neuroinflammatory phenotype.

## Discussion

Here, we have shown that the *PDGFRβ-CreER^T2^* mouse line is an effective model for selectively labelling and ablating pericytes in the CNS when crossed with *Rosa26-YFP* and *Rosa26-DTA* mice, respectively. Previous studies have achieved ∼50-60% CNS pericyte ablation using both *Pdgfrb-Flp; Cspg4-FSF-CreER :: iDTR* (cortex and hippocampus) (Nikolakopoulou et al., 2019) and *Pdgfrb-CreER^T2^ :: iDTR* (retina) transgenic mice (Eilken et al., 2017). Others have achieved ∼50% brain pericyte ablation (Choe et al., 2022), and >90% retinal pericyte ablation (Park et al., 2017) using *PDGFRβ-CreER^T2^ :: Rosa26-DTA* transgenic mice. We have identified specific Tx dosing regimens that can be used to achieve a target level of brain pericyte ablation in *PDGFRβ-CreER^T2^ :: Rosa26-DTA* transgenic mice, between ∼50% and >90% ablation. We also show that these mice can be used to achieve >90% pericyte ablation in the retina and spinal cord. By controlling the degree of pericyte ablation, it may be possible to investigate the impact that ∼50% pericyte loss, reflecting the loss reported in Alzheimer’s disease (Sengillo et al., 2013), or >80% pericyte loss, reflecting the loss reported in the core of a stroke lesion (Fernández-Klett et al., 2012), has on brain health.

Few studies have investigated the effect of pericyte loss on mouse behaviour. Previous studies reported that performance in the novel object recognition test and fear conditioning were impaired within 15 days of pericyte ablation (Nikolakopoulou et al., 2019). We report that pericyte-ablated mice displayed significant hypoactivity. This behavioural phenotype was present even after mild pericyte loss. This hypoactivity could be the result of a brain energy deficit, due to the loss of pericyte regulation in the neurovascular system (Hariharan et al., 2022). In addition to this reduction in motor activity, we showed that pericyte-ablated mice had poorer motor coordination on a beam walk task, which was only noticeable at higher levels of pericyte ablation. Furthermore, we found that the motor impairment was not due to poor muscle function, as the muscular strength of these mice was unchanged. These data may suggest that the motor impairment is centrally driven. Interestingly, we observed a reduction in motor activity when we ablated both >90% and ∼50% of pericytes, however, we only observed gliosis and BBB-impairment when ablating >90% of pericytes. This suggests that the pericyte loss itself, or pathological consequences that we did not detect, could be driving these motor phenotypes.

While pericytes are crucial for supporting vascular function in the brain, they are not significant determinants of acute vessel structure, which was unchanged in pericyte ablated mice – a finding that is consistent with previous reports (Choe et al., 2022, Nikolakopoulou et al., 2019). However, we did observe an increase in lumen area, which was accompanied by increased hypoxia. Hypoxia is increased following global (Choe et al., 2022) or focal pericyte ablation, as a study ablating a few pericytes showed that this was sufficient to increase local lumen diameter and dysregulate blood flow (Berthiaume et al., 2022). In DTA mice, pericyte ablation may drive reduced oxygen availability due to a poor distribution of blood flow, leading to a poor allocation of oxygen throughout the brain parenchyma. We also detected increased BBB leakage after pericyte ablation, which was consistent with previous acute pericyte ablation studies (Nikolakopoulou et al., 2019) but conflicts with other pericyte ablation studies (Park et al., 2017, Choe et al., 2022). We believe this is due to the different levels of pericyte ablation achieved, and the myriad of techniques with altered sensitivity to detect leakage of different sized proteins.

Reactive gliosis is one of the first cellular changes detected as the brain responds to damage (Burda and Sofroniew, 2014). We have shown that pericyte ablation leads to a widespread increase in both microglial and astrocytic reactivity, as identified by an increased area covered by IBA1^+^ microglia and GFAP^+^ astrocytes. This is consistent with other reports of astrogliosis in response to pericyte ablation (Choe et al., 2022). We also observed microglia exhibiting an abnormal clustering morphology. Similar observations have been observed during the formation of amyloid beta plaques in Alzheimer’s disease (Paasila et al., 2020), and normal-appearing white matter in multiple sclerosis (van Horssen et al., 2012), but whether these microglia clusters are protective, pathogenic, or both, remains to be determined. The severe level of glial reactivity could reflect an important role for pericytes in maintaining microglial and astrocytic homeostasis in the CNS. Alternatively, this could be a consequence of BBB leakage, or it may highlight the importance of pericytes in regulating capillary blood flow which, if disrupted, can lead to reduced oxygen availability and metabolic stress, stimulating glial reactivity in the brain. Lastly, it could also represent a neuroinflammatory phenotype. Pericytes have a well-established role in modulating neuroinflammation (Rustenhoven et al., 2017). We observed an increase in ICAM-1, a cell surface glycoprotein and leukocyte adhesion receptor (Bui et al., 2020), expression on blood vessels post-pericyte ablation, consistent with previous findings (Choe et al., 2022, Torok et al., 2021). This may indicate that pericyte loss promotes leukocyte recruitment into the brain promoting neuroinflammation.

Given the breadth of motor changes, neuroinflammation, glial activation and vascular dysfunction following pericyte ablation, these effects are consistent with a fatigue-like phenotype. Mice with fatigue-like phenotypes can experience motor impairments (Zhang et al., 2016, Lee et al., 2020) and poor motor function has been described as a clinical feature of ME/CFS (myalgic encephalomyelitis/chronic fatigue syndrome) (Lawrie et al., 2000, Siemionow et al., 2004). Additionally, chronic fatigue is a common feature of neurodegenerative diseases like stroke and multiple sclerosis, for which the fatigue is a poorly understood component of each condition with indeterminate aetiology (Christensen et al., 2008, Minden et al., 2006). People living with ME/CFS experience extreme fatigue lasting longer than 6 months which is not alleviated by rest, and is often coupled with decreased cognition, muscle weakness, headaches and pain (Bested and Marshall, 2015). Pathophysiological mechanisms thought to contribute to ME/CFS include immune, oxidative, mitochondrial, neuronal, glial, and vascular dysfunction (Cortes Rivera et al., 2019). People with ME/CFS experience reduced cerebral blood flow and poor neurovascular coupling (Yoshiuchi et al., 2006, Biswal et al., 2011, Shan et al., 2020). These effects are similar to those displayed following pericyte ablation, suggesting that pericyte dysfunction could drive the microvascular impairment and neuroinflammatory response in people living with ME/CFS.

*Pdgfrβ* is expressed by cells other than pericytes in the CNS including VSMCs and perivascular fibroblasts (Vanlandewijck et al., 2018, Dorrier et al., 2022), and by pericytes and other mesenchymal-derived cells outside of the CNS (Baek et al., 2022, Cornuault et al., 2023, Wang et al., 2018). We showed that DTA activation did not deplete VSMCs, which may be due to the 6-fold higher expression of PDGFRβ protein and 4-fold higher mRNA levels in pericytes compared to VSMCs (Nikolakopoulou et al., 2019, Vanlandewijck et al., 2018). However, further refinement, such as use of recently identified promoters like *Atp13a5*, which are more unique to CNS pericytes (Guo et al., 2021), or the organ specific administration of 4-hydroxytamoxifen, could be leveraged to specifically target brain pericytes in future work. Both male and female mice were used for this study. Sex was not considered as a variable throughout the analyses, consistent with previous pericyte ablation studies (Eilken et al., 2017, Nikolakopoulou et al., 2019, Kisler et al., 2020, Choe et al., 2022).

## Conclusions

Here, we showed that the *PDGFRβ-CreER^T2^* mouse line, when crossed with *Rosa26-YFP* or *Rosa26-DTA* mice, provides an effective model for labelling or ablating brain pericytes. We demonstrated that pericyte ablation induces behavioural changes in mice, inducing a fatigue-like phenotype, as evidenced by hypoactivity and impaired motor coordination. Ablating over 80% of brain pericytes resulted in vascular changes including increased lumen area and BBB leakage, while leaving vessel structure unchanged. Pericyte ablation also led to significant gliosis in the brain, as indicated by microgliosis and astrogliosis. These consequences of pericyte ablation may be due to impaired energy supply in the brain, which is a leading hypothesis of ME/CFS causation. These findings reveal that pericytes are vital for brain function and warrant further investigation into the role of pericytes in cerebrovascular regulation, neuroinflammation and motor function.

## Statements and Declarations

### Author contributions

**JaMC:** conceptualisation, data curation, formal analysis, investigation, methodology, software, validation, visualisation, writing – original draft, writing – review & editing. **LSB:** investigation, methodology, writing – review & editing. **GPM:** investigation, methodology, software, supervision, writing – review & editing. **AJF:** conceptualisation, formal analysis, investigation, methodology, software, writing – review & editing. **JoMC:** formal analysis, methodology, resources, software, writing – review & editing. **CLC:** methodology, supervision, writing – review & editing. **KMY:** conceptualisation, funding acquisition, investigation, methodology, resources, project administration, supervision, visualisation, writing – original draft, writing – review & editing. **BAS:** conceptualisation, funding acquisition, investigation, methodology, resources, project administration, supervision, visualisation, writing – original draft, writing – review & editing.

## Acknowledgements

We’d like to thank Loic Auderset and Kalina Makowiecki for support with oral gavage procedures. We’d like to thank the Animal Services Team at the University of Tasmania, especially Ellen Bennett and Peta Yates for their assistance with animal care and management.

## Data availability

The image files and data described in this manuscript can be obtained from the corresponding author by reasonable request.

## Ethics approval

All animal experiments were carried out on project A0018606, which was approved by the University of Tasmania’s Animal Ethics Committee.

## Consent for publication

This publication does not include data from human research participants.

