## Supplementary material for "Pericyte ablation causes hypoactivity and reactive gliosis in adult mice"

### Supplementary methods

#### Pericyte and OPC DsRed density

The density and proportion of pericytes and OPCs labelled with DsRed was manually determined as previously described (Morris et al., 2023) with minor changes. Images were opened and blinded in QuPath (Bankhead et al., 2017) and regions of interest (ROIs) determined using the Allen Brain Atlas (Lein et al., 2007). One ROI was analysed per hemisphere from one brain section per mouse. $0.5mm^2$ ROIs were placed within the somatosensory cortex (spanning layers 2/3 to 6) and thalamus. Within the hippocampus, ROIs were manually traced around Cornu Ammonis (CA)2 and CA3. The corpus callosum ROI was traced from the dorsal-ventral midline to the lateral ventricle. A low threshold cell detection was run using the DAPI channel to detect all nucleated cells. Pericytes were identified morphologically (Attwell et al., 2016) and OPCs were identified with morphology and PDGFRα immunoreactivity.

#### Basement membrane length

To determine basement membrane length, immunofluorescent images of tomato lectin labelling from the somatosensory cortex and thalamus ($0.65mm^2$, one region per section, one section per mouse) were blinded and analysed in FIJI-imageJ. The segmented line tool was used to trace along the length of positive signal in each image. ROI manager was then used to extract the total length annotated from each image.

#### Somatostatin$^+$ cell density

For determination of somatostatin (SST) interneuron density, SST$^+$ cells were manually counted using QuPath. Images were blinded and a ROI was drawn in the hippocampus (entire hippocampus, one section per mouse). A low threshold cell detection was run using the DAPI channel to detect all nucleated cells, and $SST^+$ cells were manually counted.

**Supplementary Table 1. Health score parameters used for monitoring the health of mice.**

| Score Category | Score Criteria |
|---|---|
| Coat | Healthy = 0<br>Rough, soiled = 1 |
| Eyes (and other components of the mouse grimace score that can indicate pain) | Normal = 0<br>Moderate i.e. eyes taught, ears slightly back = 1<br>Severe i.e. eyes closed, ears back = 2 |
| Respiration | Normal = 0<br>Rapid, shallow = 1<br>Deep / laboured, noisy = 2 |
| Social parameters | Normal = 0<br>Not grooming or nesting, isolated from cage mates, shows signs of compulsive behaviour = 1<br>Highly docile / aggressive, vocal, resistant to handling = 2 |
| Posture | Normal = 0<br>Hunched = 1 |
| Movement within cage | Normal = 0<br>Mild lassitude, needs prompting to move around home cage, but can move relatively unhindered i.e. can access food and water = 0.5<br>Needs prompting to move around home cage and resulting movement is slower or less co-ordinated = 1<br>No movement when prompted = 2 |
| Movement outside of cage (on a flat surface) | Normal = 0<br>Needs prompting = 1<br>No movement when prompted = 2 |
| Gait | Normal = 0<br>Wide hindlimb stance = 0.5<br>Slight tremor when walking or moves freely with slight favouring of particular paws = 1<br>Dragging belly or hindlimbs, forelimb placement errors or severe tremor when walking = 2 |

**Supplementary Table 2. Antibodies and fluorescent labels**

| Antibody/stain | Abbreviation | Host species | Manufacturer | Cat. number | RRID | Dilution |
|---|---|---|---|---|---|---|
| Donkey anti-Goat IgG (H+L) Cross-Adsorbed Secondary Antibody, Alexa Fluor™ 647 | - | Donkey | Invitrogen | A-21447 | AB_141844 | 1:1000 |
| Donkey anti-Mouse IgG (H+L) Alexa Fluor™ 647 | - | Donkey | Invitrogen | A-31571 | AB_162542 | 1:1000 |
| Donkey anti-Rabbit IgG (H+L) Highly Cross-Adsorbed Secondary Antibody, Alexa Fluor™ 594 | - | Donkey | Invitrogen | A-21207 | AB_141637 | 1:1000 |
| Donkey anti-Rat IgG (H+L) Alexa Fluor™ 488 | - | Donkey | Invitrogen | A-21208 | AB_2535794 | 1:1000 |
| Donkey anti-Rat IgG (H+L) Alexa Fluor™ 594 | - | Donkey | Invitrogen | A-21209 | AB_2535795 | 1:1000 |
| Donkey anti-Rat IgG (H+L) Alexa Fluor™ 594 | - | Donkey | Invitrogen | A-21209 | AB_2535795 | 1:1000 |
| Goat anti-Guinea Pig IgG (H+L) Highly Cross-Adsorbed Secondary Antibody, Alexa Fluor™ 647 | - | Goat | Invitrogen | A-21450 | AB_141882 | 1:1000 |
| Polyclonal Rabbit Anti-Human Albumin (Multipurpose) | anti-albumin | Rabbit | Dako | A009002 | AB_2943648 | 1:500 |
| Purified Rat anti-mouse CD31 Antibody | anti-CD31 | Rat | Biolegend | 102501 | AB_312908 | 1:500 |

| | | | | | | |
|---|---|---|---|---|---|---|
| Anti-GFAP antibody, goat polyclonal | anti-GFAP | Goat | Abcam | ab53554 | AB_880202 | 1:2000 |
| Hypoxyprobe Plus Kit - Mouse FITC-MAb plus rabbit anti-FITC HRP | anti-HypoxyProbe | Mouse | Hypoxyprobe | HP2-100Kit | AB_2811309 | 1:100 |
| Anti-Iba1 Rabbit antibody | anti-IBA1 | Rabbit | Wako | 019-19741 | AB_839504 | 1:2000 |
| Anti-ICAM1 antibody [YN1/1.7.4] | Anti-ICAM | Rat | Abcam | ab119871 | AB_10900211 | 1:200 |
| Recombinant Anti-Myelin Basic Protein antibody | anti-MBP | Rat | Abcam | ab7349 | AB_305869 | 1:100 |
| Anti-PDGFR alpha Antibody | anti-PDGFRα | Goat | R&D Systems | AF1062 | AB_2236897 | 1:500 |
| CD140b (PDGFRB) Monoclonal Antibody (APB5), eBioscience™ | anti-PDGFRβ | Rat | Invitrogen | 14-1402-82 | AB_467493 | 1:200 |
| Recombinant Anti-Somatostatin 28 antibody | anti-somatostatin | Rabbit | Abcam | ab111912 | AB_10903864 | 1:250 |
| GFP Polyclonal Antibody | anti-YFP | Rabbit | Invitrogen | A-11122 | AB_221569 | 1:500 |
| Anti-alpha smooth muscle Actin antibody | anti-αSMA | Rabbit | Abcam | ab5694 | AB_2223021 | 1:1000 |

| | | | | | | |
|---|---|---|---|---|---|---|
| DAPI (4',6-Diamidino-2-Phenylindole, Dihydrochloride) | DAPI | - | ThermoFisher | D1306 | - | 1:10,000 |
| Lectin from Lycopersicon esculentum (tomato) | Lectin | - | Sigma-Aldrich | L0401-1MG | - | 1:500 |

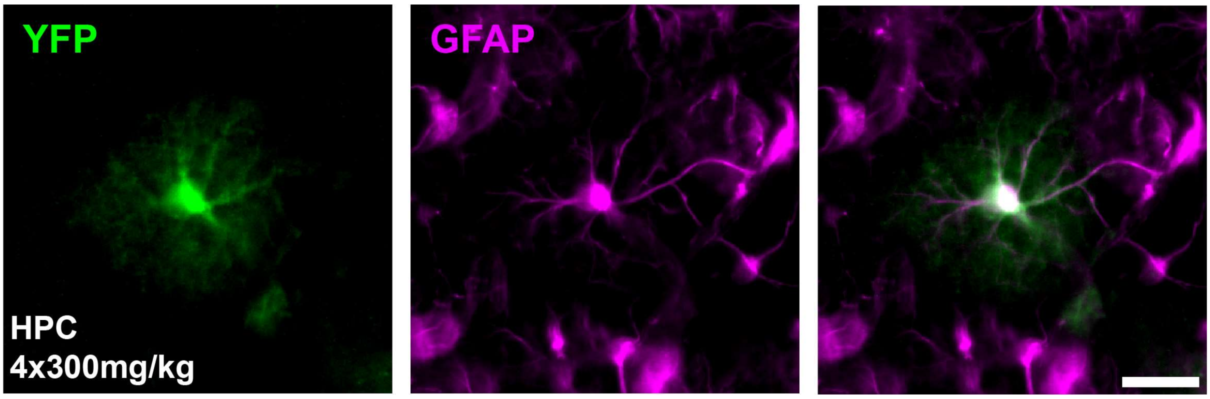


**Supplementary Figure 1. A subpopulation of astrocytes express YFP in transgenic mice 7-days post-tamoxifen administration.** A hippocampal (HPC) YFP (yellow fluorescent protein) positive astrocyte co-labelled with GFAP (glial fibrillary acidic protein). Scale = 20µm.

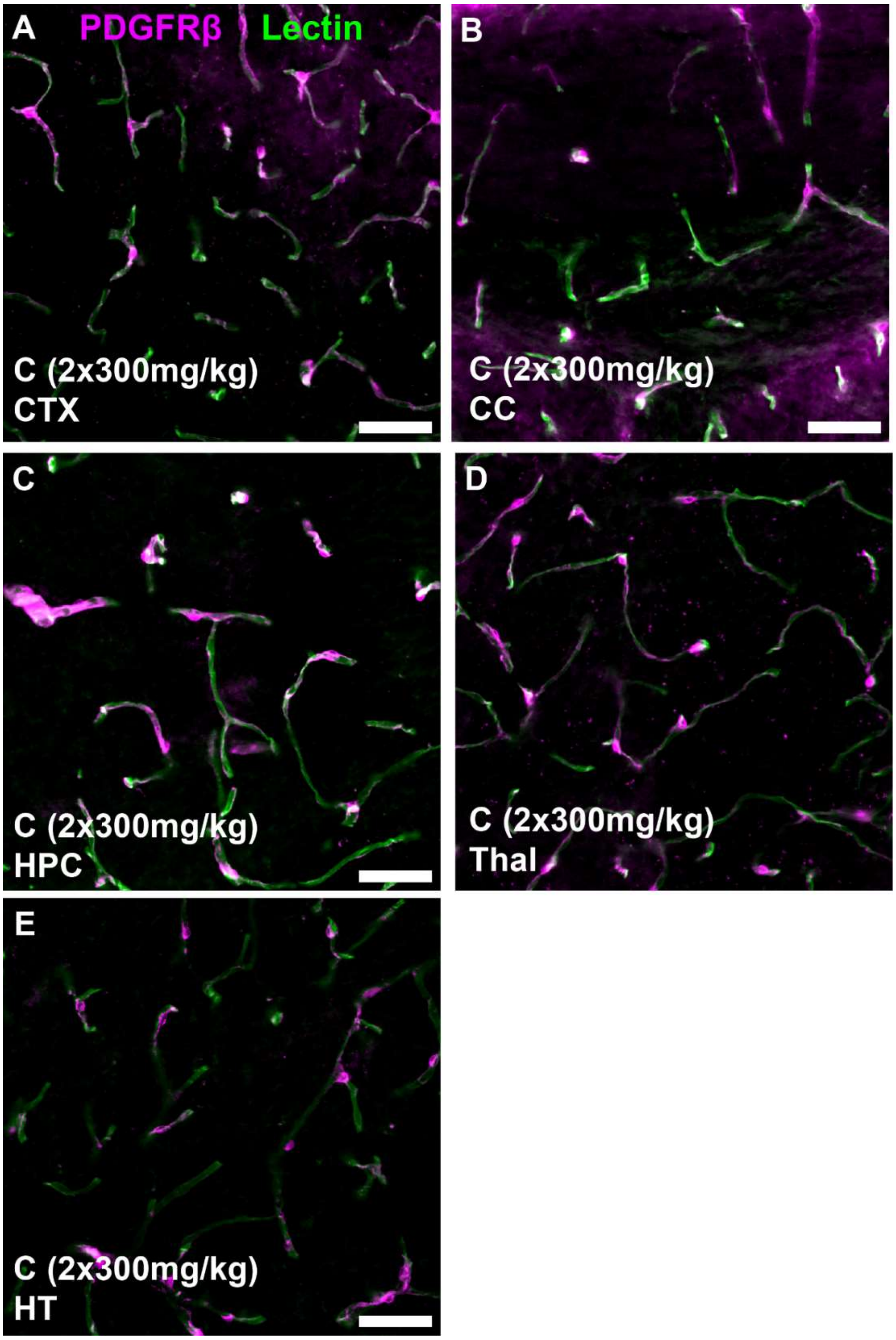


**Supplementary Figure 2. Pericytes express PDGFRβ throughout the brain. A-E)** Immunofluorescent images of PDGFRβ (platelet-derived growth factor receptor beta) labelling of pericytes (magenta) and tomato lectin labelling of blood vessels (green) in the somatosensory cortex (CTX, A), corpus callosum (CC, B), hippocampus (HPC, C), thalamus (Thal, D) and hypothalamus (HT, E) of control mice. Scale = 50µm.

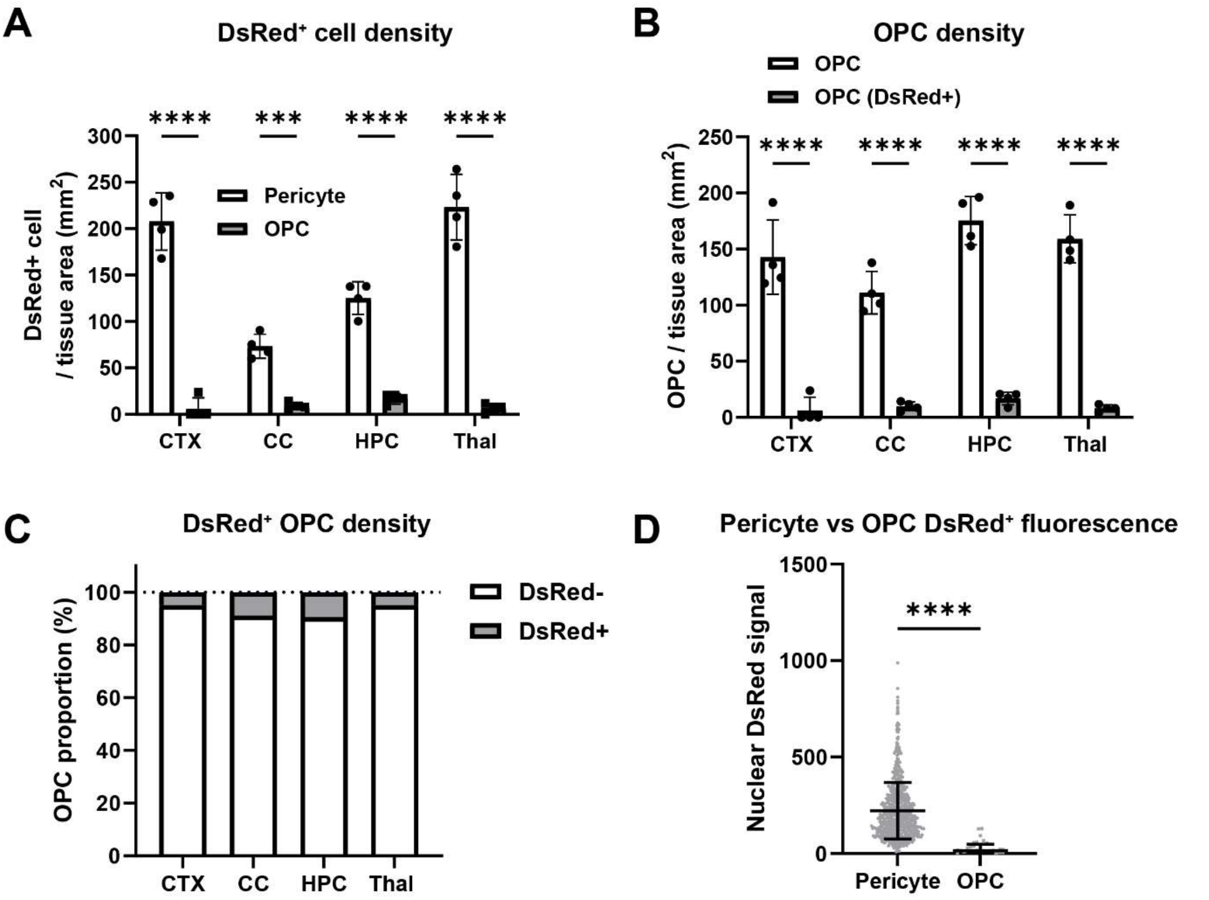


**Supplementary Figure 3. Oligodendrocyte progenitor cells rarely label with detectable levels of DsRed in the brains of NG2DsRed mice. A)** Density of DsRed$^+$ pericytes and OPCs in the somatosensory cortex (CTX), corpus callosum (CC), hippocampus (HPC), thalamus (Thal). Two-way ANOVA: brain region: F (3, 24) = 25.70, $p < 0.0001$; cell type: F (1, 24) = 481.9, $p <0.0001$; interaction: F (3, 24) = 29.65, $p < 0.0001$. **B)** Density of DsRed$^+$ and DsRed- OPCs in the CTX, CC, HPC and thal. Two-way ANOVA: brain region: F (3, 24) = 5.569, $p = 0.0048$; cell type: F (1, 24) = 465.5, $p < 0.0001$; interaction: F (3, 24) = 4.033, $p = 0.0186$. **C)** Proportion of OPCs that are DsRed- vs DsRed$^+$ in the CTX, CC, HPC and thal. **D)** Mean nuclear DsRed signal from pericytes and OPCs across all brain regions (CTX, CC, HPC, Thal) quantified. Mann-Whitney test: $p < 0.0001$. Data shown as mean ± SD, with individual data points for each animal. Two-way ANOVA with Šidák's multiple comparisons test and Mann-Whitney tests. *** p < 0.001, **** p < 0.0001.

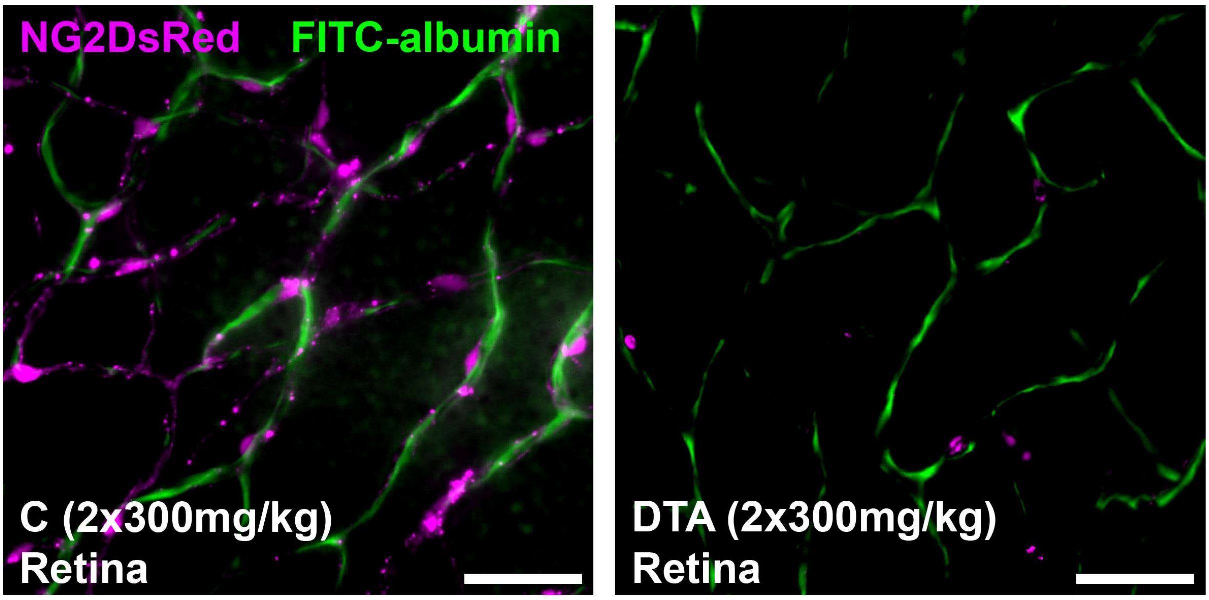


**Supplementary Figure 4. Representative image of retinal pericytes in transgenic mice 7-days following tamoxifen administration.** Fluorescent images of retinal pericytes (NG2DsRed, magenta) on capillaries (FITC-albumin, green) in control and DTA mice. Scale = 40µm.

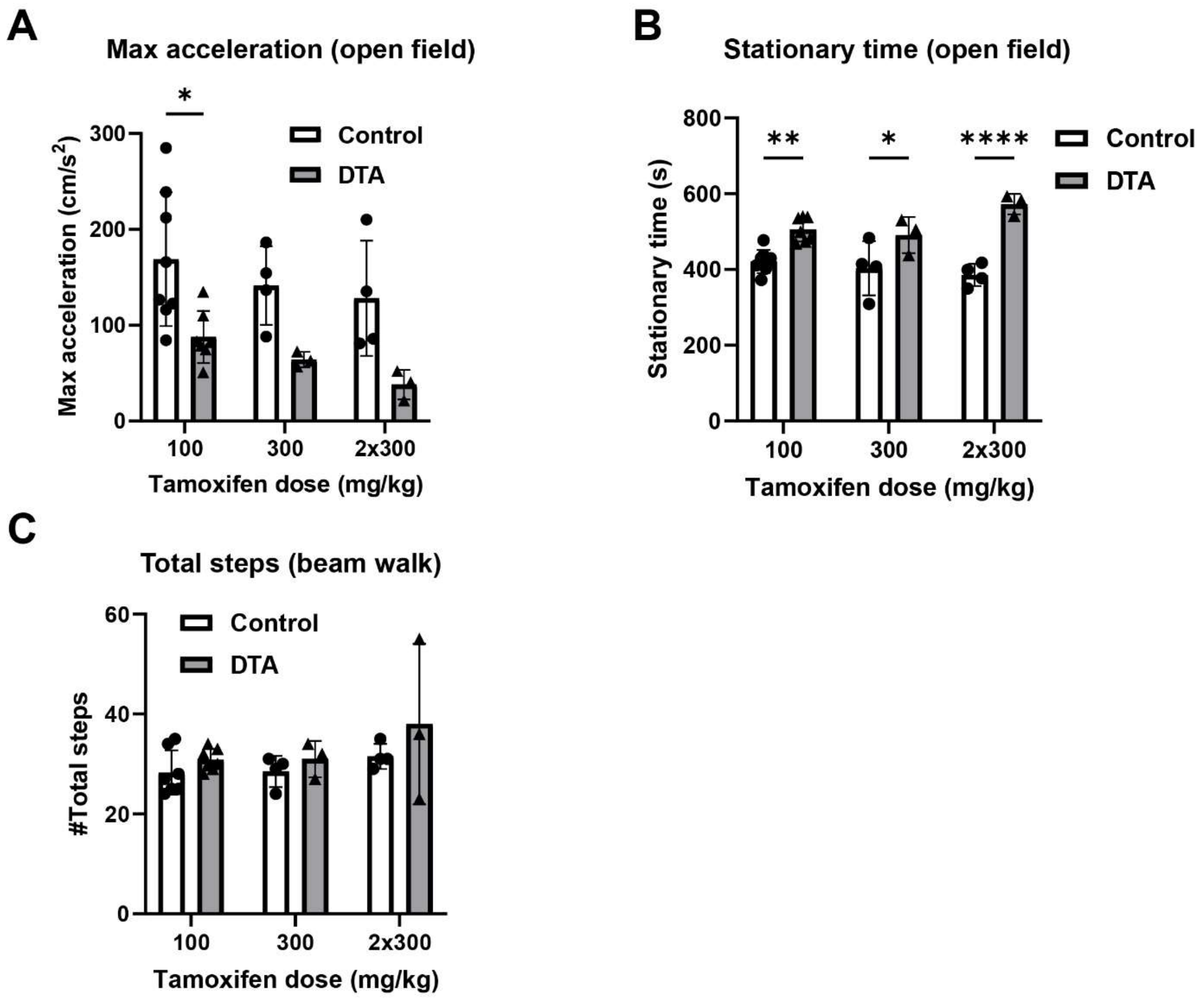


**Supplementary Figure 5. Pericyte ablation reduces max acceleration, increases time spent stationary and time to cross the beam walk. A-C)** Max acceleration (A; two-way ANOVA: tamoxifen dose: F (2, 23) = 2.150, $p$ = 0.1393; genotype: F (1, 23) = 17.98, $p$ = 0.0003; interaction: F (2, 23) = 0.03165, $p$ = 0.9689) and time spent stationary (B; two-way ANOVA: tamoxifen dose: F (2, 23) = 1.102, $p$ = 0.3490; genotype: F (1, 23) = 56.23, $p$ < 0.0001; interaction: F (2, 23) = 4.172, $p$ = 0.0284) in the open field, and total steps to cross the beam walk (C) in control and DTA mice. Data shown as mean ± SD, with individual data points for each animal. Two-way ANOVA with Šidák's multiple comparisons test. * p < 0.05, ** p < 0.01, **** p < 0.0001.

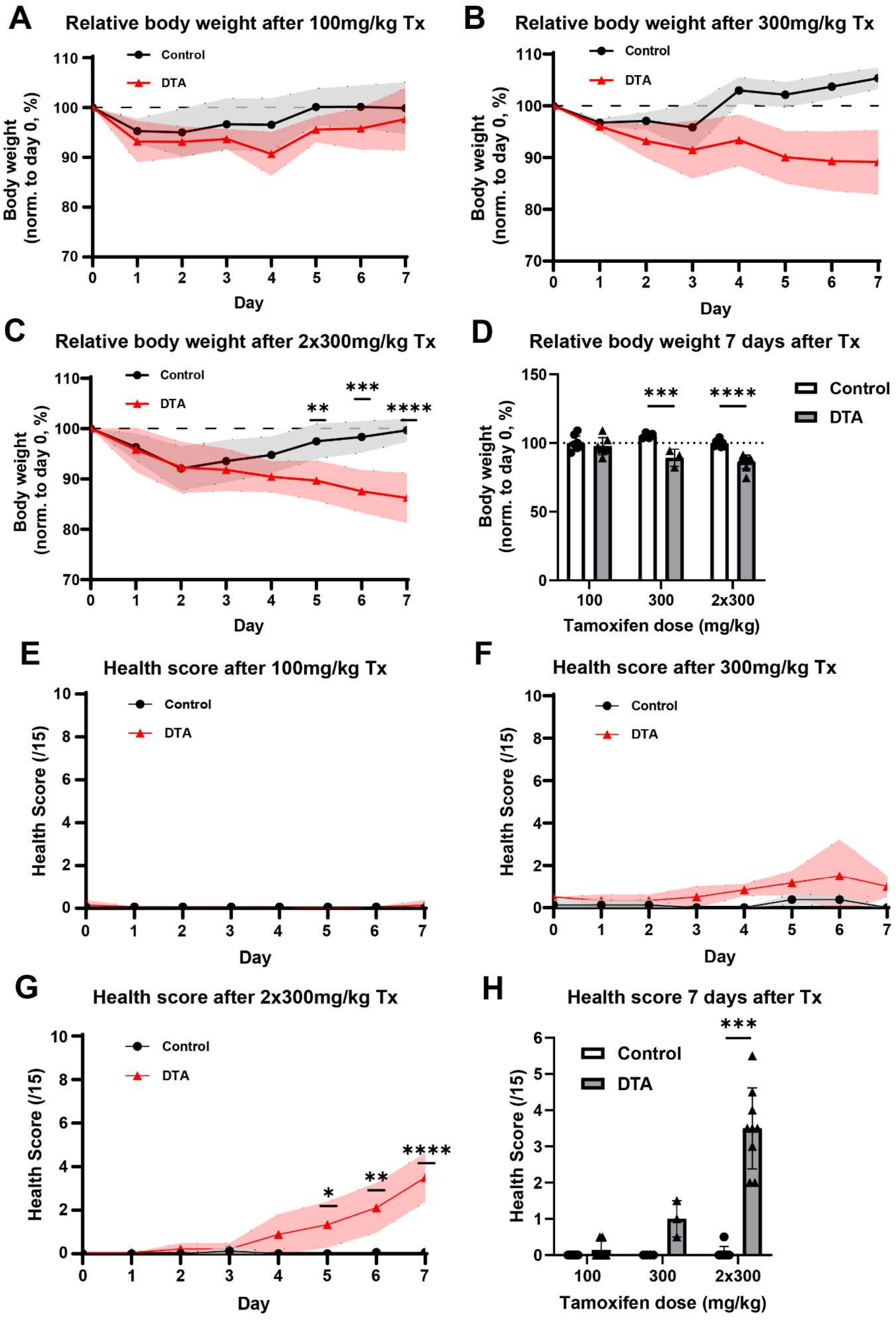
A
Relative body weight after 100mg/kg Tx
Body weight (norm. to day 0, %)
Control
DTA
Day
B
Relative body weight after 300mg/kg Tx
Body weight (norm. to day 0, %)
Control
DTA
Day
C
Relative body weight after 2x300mg/kg Tx
Body weight (norm. to day 0, %)
Control
DTA
Day
D
Relative body weight 7 days after Tx
Body weight (norm. to day 0, %)
Control
DTA
Tamoxifen dose (mg/kg)
E
Health score after 100mg/kg Tx
Health Score (/15)
Control
DTA
Day
F
Health score after 300mg/kg Tx
Health Score (/15)
Control
DTA
Day
G
Health score after 2x300mg/kg Tx
Health Score (/15)
Control
DTA
Day
H
Health score 7 days after Tx
Health Score (/15)
Control
DTA
Tamoxifen dose (mg/kg)

**Supplementary Figure 6. βCre-DTA mice experience significant weight loss and increased health score when dosed with ≥300mg/kg tamoxifen. A-C)** Normalised weight of control and DTA mice over 7 days when dosed with 100mg/kg (A), 300mg/kg (B) or two consecutive daily doses of 300mg/kg (2x300mg/kg; C; two-way ANOVA: time: F (3.493, 52.39) = 23.39, $p < 0.0001$; genotype: F (1, 15) = 10.61, $p = 0.0053$; interaction: F (7, 105) = 23.10, $p < 0.0001$) tamoxifen (Tx). **D)** Normalised weight of control and DTA mice 7-days after varying doses of tamoxifen. Two-way ANOVA: tamoxifen dose: F (2, 33) = 6.197, $p = 0.0052$; genotype: F (1, 33) = 40.94, $p < 0.0001$; interaction: F (2, 33) = 7.468, $p = 0.0021$. **E-G)** Health scores (higher=worse) of control and DTA mice over 7 days when dosed with 100mg/kg (E), 300mg/kg (F) or 2x300mg/kg (G; two-way ANOVA: time: F (3.197, 47.96) = 23.01, $p < 0.0001$; genotype: F (1, 15) = 60.21, $p < 0.0001$; interaction: F (7, 105) = 22.02, $p < 0.0001$) tamoxifen. **H)** Health scores of control and DTA mice 7-days after varying doses of tamoxifen. Mann-Whitney test: 100mg/kg, $p = 0.2$; 300mg/kg, $p = 0.056327$, 2x300mg/kg, $p = 0.000123$. Data shown as mean ± SD, with individual data points for each animal. Two-way ANOVA with Šidák's multiple comparisons test. * p < 0.05, ** p < 0.01, *** p < 0.001, **** p < 0.0001.

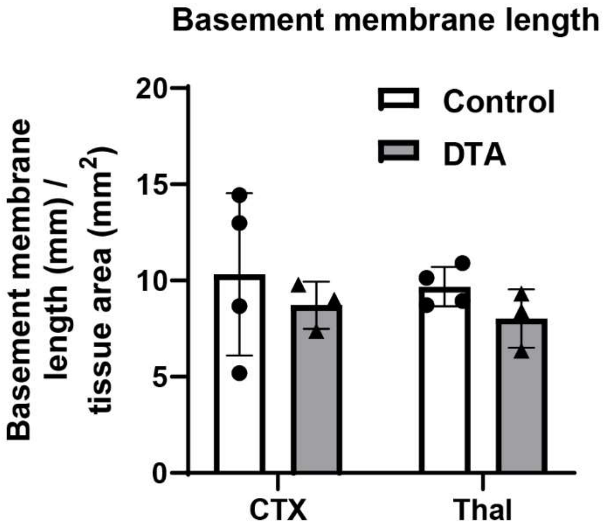


**Supplementary Figure 7. Basement membrane length remains unchanged 7-days post-pericyte ablation.** Basement membrane length of control and DTA mice dosed with two consecutive daily doses of 300mg/kg tamoxifen. CTX = somatosensory cortex, Thal = thalamus. Data shown as mean ± SD, with individual data points for each animal.

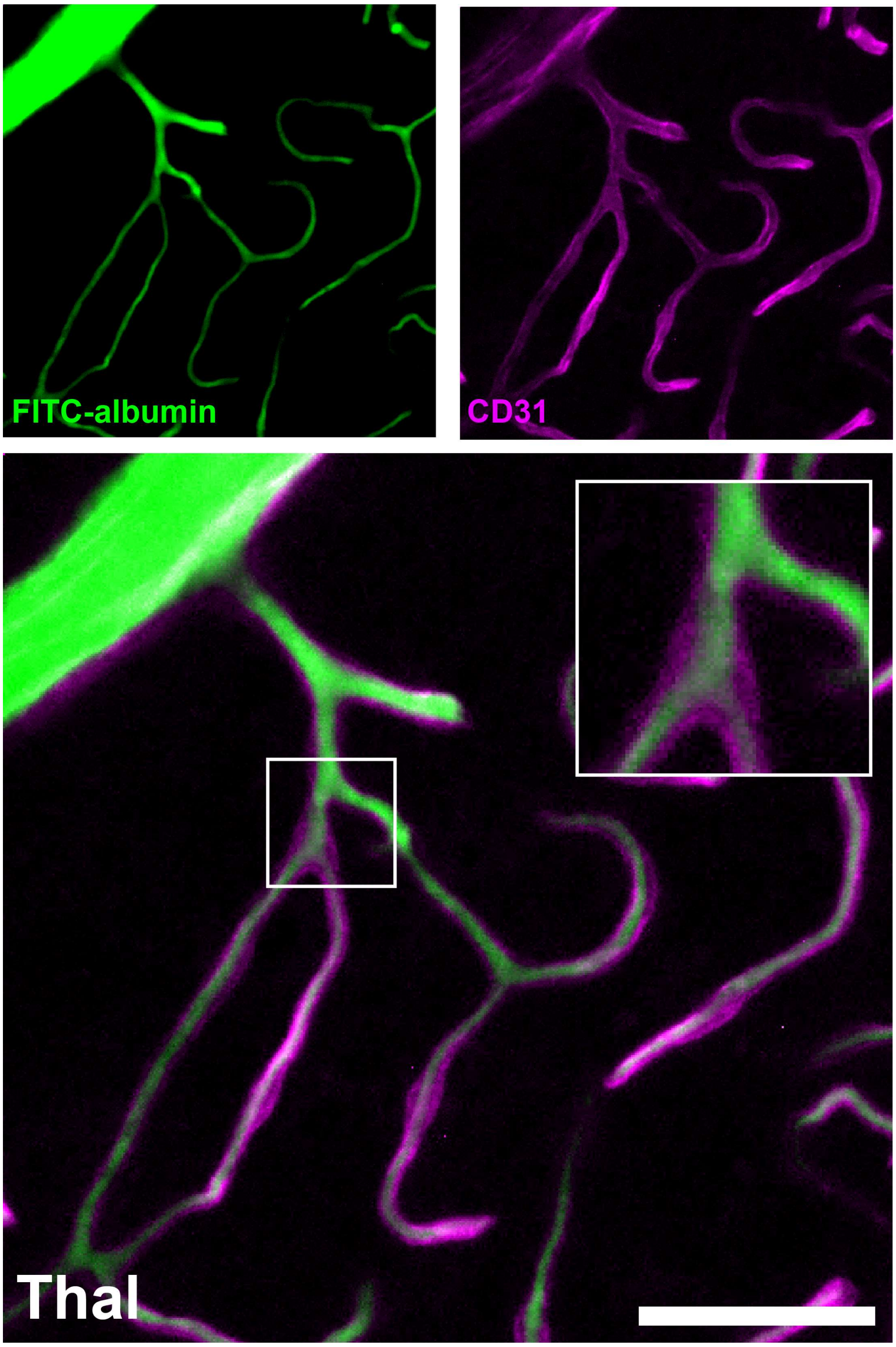


**Supplementary Figure 8. Representative image of FITC-albumin gelatin cast within $CD31^+$ endothelial cells of capillaries.** FITC-albumin (green) labels the vessel lumen. CD31 immunolabelling (magenta) identifies the endothelial cell layer of blood vessels. Thal = thalamus. Scale = 40µm.

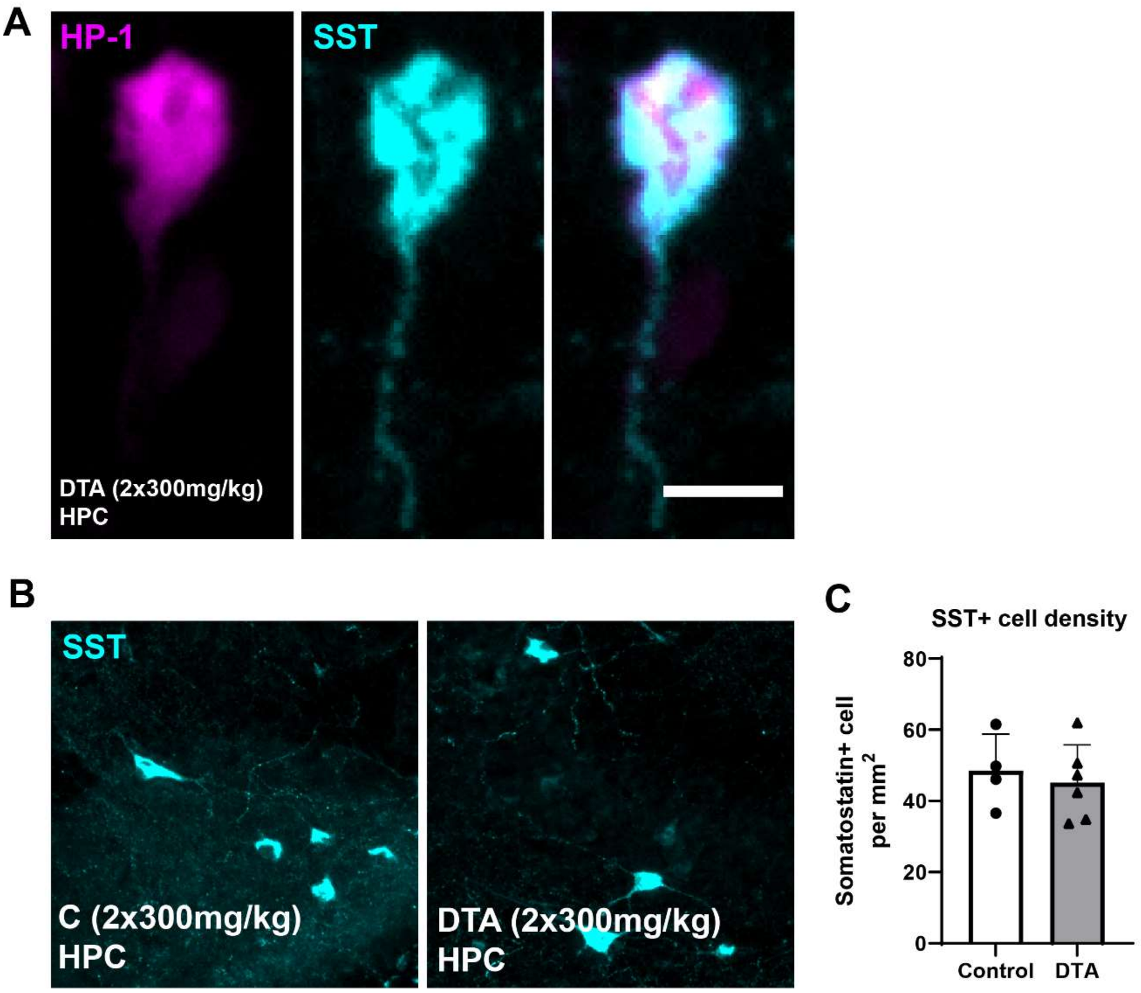


**Supplementary Figure 9. Somatostatin positive interneurons label with hypoxyprobe-1. A)** Immunofluorescent labelling of a somatostatin $(SST)^{+}$ interneuron co-labelled with hypoxyprobe-1 in a βCre-DTA mouse. Scale = 10µm. **B)** Representative images of SST immunolabelling in control and DTA mice. Scale = 40µm. **C)** $SST^{+}$ cell density in the hippocampus of control and DTA mice. Data shown as mean ± SD, with individual data points for each animal.

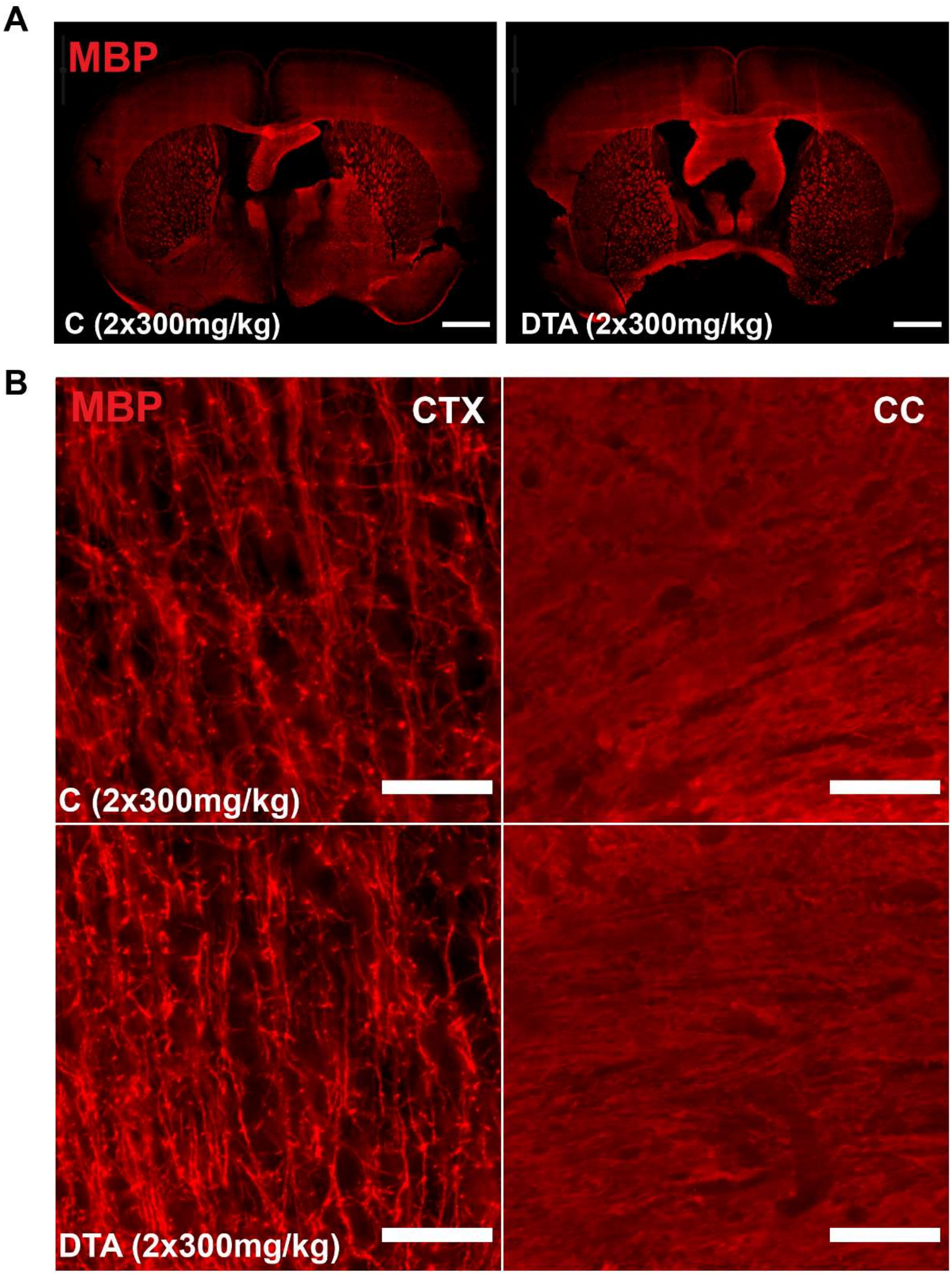


**Supplementary Figure 10. Gross myelin structure is unchanged 7-days post-pericyte ablation. A)** Full coronal brain images of myelin basic protein (MBP) immunolabelling in control and DTA mice. **B)** Immunofluorescent images of MBP labelling in the somatosensory cortex (CTX) and corpus callosum (CC) of control and DTA mice. Scale = 40µm.
